# Fragment-Based Development of NSP14 Exonuclease Inhibitors Confounded by Batch-to-Batch Variability

**DOI:** 10.1101/2025.11.21.689747

**Authors:** Jesse A. Coker, Rong Sun, Paul M. Polzer, Todd Romigh, Christopher M. Goins, Nancy S. Wang, Jae U. Jung, Shaun R. Stauffer

## Abstract

Point mutations in the exonuclease (ExoN) site of non-structural protein 14 (NSP14) compromise the fitness of betacoronaviruses like SARS-CoV-2, implicating NSP14 ExoN inhibition as an antiviral strategy. However, there are no advanced compounds that inhibit NSP14’s ExoN activity. Building upon the reported crystal structures of two fragments bound to NSP14’s ExoN site, we identified a series of 3,5-disubsituted pyrazoles that bound to and inhibited NSP14 ExoN. However, upon resynthesis, we discovered that these putative leads were false positives, perhaps due to contaminating divalent cations which potently inhibit NSP14 ExoN. Our results provide a cautionary tale to the field about the sensitivity of NSP14 to divalent cations and illustrate the challenges associated with directly targeting the NSP14 ExoN site via fragment merging.

## MAIN TEXT

Despite the remarkable success of vaccines against SARS-CoV-2, COVID-19 remains an endemic threat that kills hundreds of people daily in the US.^1^ Viral RNA has been detected in >50% of long COVID patients, and early evidence suggests that antiviral treatments can alleviate these symptoms,^2-4^ emphasizing the important role of antivirals in the fight against pandemic betacoronaviruses. To date, however, there are only three FDA-approved SARS-CoV-2 antivirals: two targeting the RNA-dependent RNA polymerase (molnupiravir and remdesivir), and one targeting the main protease (nirmatrelvir). It is critical to diversify the available SARS-CoV-2 antivirals, especially against novel targets.

Non-structural protein 14 (NSP14) is the main 3’ to 5’ proofreading RNA exonuclease (ExoN) of SARS-CoV-2. NSP14 undergoes a profound conformational change upon binding to its obligate co-activator, NSP10, which activates RNA hydrolysis through an evolutionarily conserved protein-protein interaction (PPI) that opens the NSP14 “hand-like” ExoN domain.^5, 6^ ExoN-inactivating point mutants obliterate the replication of SARS-CoV-2, the related betacoronavirus MERS, and mouse hepatitis virus.^7-11^ The PPI between NSP14:NSP10 has also been experimentally validated as essential for viral replication.^12^ Furthermore, NSP14 ExoN activity has been linked to viral immune evasion via impairment of host protein translation.^13^ Therefore, with a dual role in viral fitness and immune escape, NSP14 ExoN represents an attractive next-generation antiviral target.

However, the compounds targeting NSP14 ExoN are much less advanced than those targeting NSP14’s secondary methyltransferase active site.^14-18^ Small biochemical screens have revealed that promiscuous compounds such as patulin, aurintricarboxylic acid, ebselen, sofalcone, and bismuth-containing salts can inhibit NSP14 ExoN, but these compounds are unsuitable for further translation. ^8, 19^ ^7, 20^

We decided to leverage structure-guided drug design to identify novel, more drug-like NSP14 ExoN inhibitors. In 2023, Imprachim *et al*. disclosed a high-throughput XChem crystallographic fragment screen against SARS-CoV-2 NSP14 that identified two fragments (**1**, PDB: 5SMD; **2**, PDB: 5SKY) bound to the NSP14 ExoN active site (**Figure 1A**).^5^ **1** and **2** bind adjacently to each other near His268, with the pyrimidine **1** binding on the “left” side of the pocket near the RNA binding groove and the pyrazole **2** binding on the “right” side of the pocket via two hydrogen bonds to Gln108 and the catalytic Asp273 (**Figure 1A**). **2** reaches rightward towards the NSP14:NSP10 PPI interface, where two disordered loops (His95—Gly102; Gly123—Ile150) must rearrange to become ordered upon binding to the co-activating NSP10.

**Figure 1.**
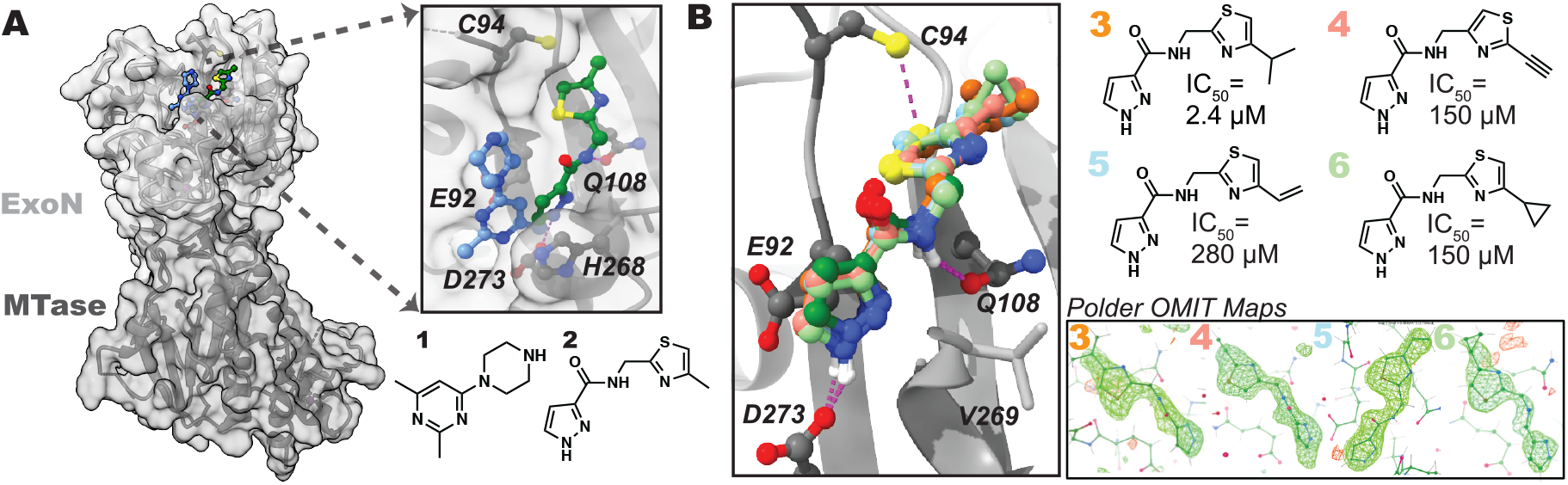
**A**. Crystal structures of NSP14 bound to **1** (blue) and **2** (green) (PDB: 5SMD, 5SKY). **B**. Crystal structures of **3** (orange), **4** (pink), **5** (blue), and **6** (light green) bound to the ExoN active site of NSP14 in the same orientation as **2** (dark green). Polder difference maps (inset; contoured at 3σ) indicate unambiguous ligand electron density (PDB: 9NHA, 9NIO, 9NAZ, 9NFP).

Because XChem screening uses “PanDDA” analysis to identify density from even weakly-bound fragments,^21^ we sought to validate the binding pose of the XChem hits by bespoke crystallography using a small set of newly synthesized analogs of **2**. Using a crystal soaking system, we solved four additional high-resolution (2.0—2.4 Å) structures of NSP14 bound to pyrazole-amides **3** (PDB: 9NHA), **4** (PDB: 9NIO), **5** (PDB: 9NAZ), and **6** (PDB: 9NFP). We observed identical binding poses, and unambiguous ligand density (confirmed by Polder OMIT maps),^22^ for all four fragments. We also validated the presence of two conserved hydrogen bonds to NSP14 (Asp273 and Gln108, **Figure 1B**).

Neither **1** (pIC_50_ = 3.4 ± 0.1, n = 4) nor **2** (pIC_50_ < 3.3, n = 5) potently inhibited NSP14 ExoN in a biochemical activity assay, while the isopropyl thiazole **3** (pIC_50_ = 5.6 ± 0.2, n = 4), the alkyne thiazole **4** (IC_50_ = 3.8 ± 0.2, n = 5), the alkene thiazole **5** (pIC_50_ = 3.6 ± 0.3, n = 5), and the cyclopropyl thiazole **6** (pIC_50_ = 4.3 ± 0.3, n = 9) were more active. To further enhance potency, we evaluated multiple **1** + **2** mergers inspired by the “ideal” merged compound suggested by the Fragmentstein algorithm (**Figure 2A**).^23^ Neither fusion of the pyrazole and pyrimidine rings into a bicyclic 7-deazapurine (such as **7**—**9**) nor direct substitution of aromatic rings at the 5-position of the pyrazole (such as **10** and **11**) enhanced activity. However, multiple homologated 5-alkyl pyrazoles (**12** and **13**) showed robust and complete ExoN inhibition (pIC_50_ = 4.3 ± 0.2, n = 3 and pIC_50_ = 4.4 ± 0.2, n = 4, respectively).

**Figure 2.**
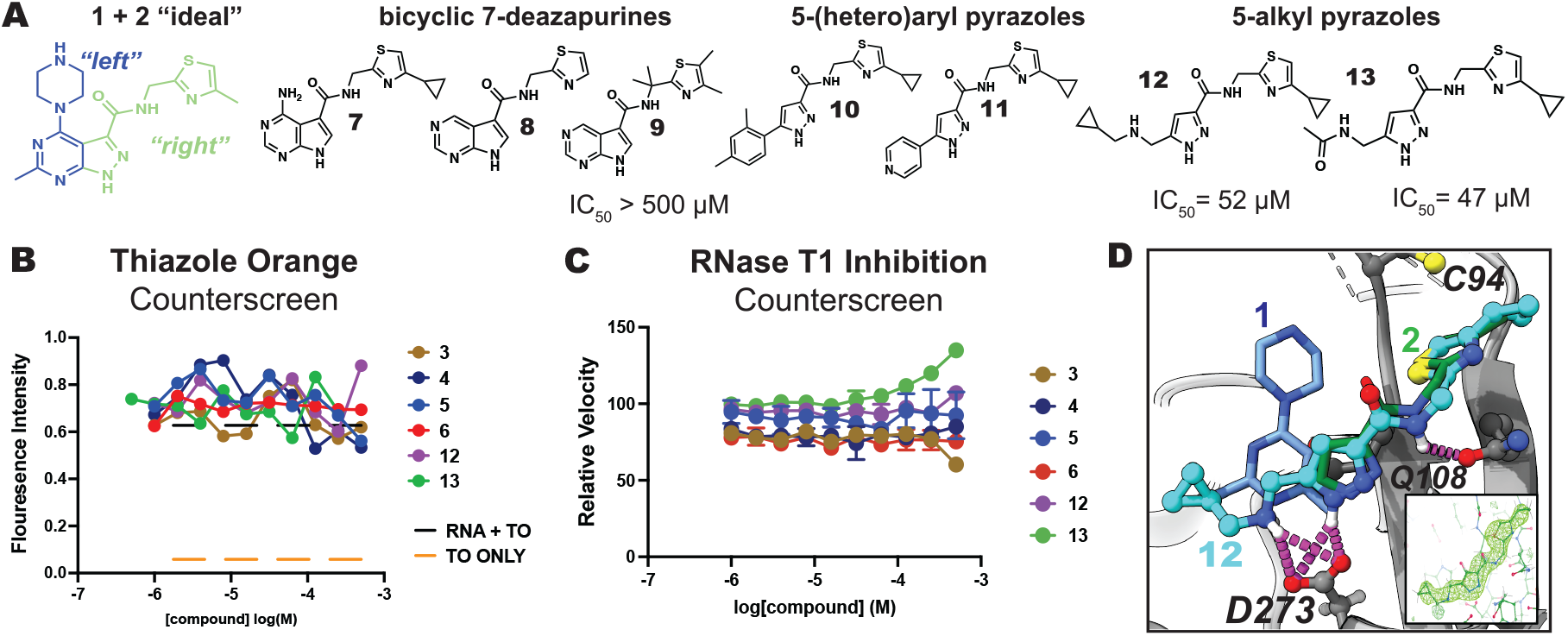
**A**. Design and testing of **1** + **2** merged compounds. **B**. Thiazole orange dsRNA intercalation assay. **C**. RNAse T1 activity assay. **D**. Crystal structure of **12** (cyan spheres; Polder map inset) bound to NSP14 (PDB: 9NHU), overlaid with **1** (blue sticks) and **2** (green sticks).

We confirmed on-target activity as recommended by others for NSP14 inhibitors by demonstrating that **12** and **13** neither bound to substrate dsRNA (by thiazole orange competition assay, **Figure 2B**), nor inhibited the unrelated endonuclease RNAse T1 (**Figure 2C**).^24^ The unmerged fragments, **3—6**, were also clean in these two counterscreens (**Figure 2B, C**). We confirmed the binding mode of merged 5-alkyl pyrazoles through a crystal structure with **12** (2.3 Å; PDB: 9NHU). **12**’s 5-cyclopropylamine substituent reaches leftward in the pocket and overlays closely with the pyrimidine N from **1**, while also contributing to a bidentate H-bond with the catalytically essential Asp273 (**Figure 2D**).^6, 9^

Despite a clean counter-screen against RNAse T1, respectable ExoN inhibition activity, and structural validation of **12**’s merged design, we remained concerned about the possibility of spurious activity due to steep SAR in the right-hand thiazole, especially the surprising low micromolar activity of the isopropyl thiazole **3** (**Figure 1**). Indeed, the isopropyl thiazole analogs of **12** and **13** (**15** and **16**) were inactive (pIC_50_ < 3.3, n = 2; **Figure 3B**). We were also troubled by the crystal structure of **14**, a 7-azaindazole merged compound that displayed weak ExoN inhibition (pIC_50_ = 3.8 ± 0.2, n = 4), which we unexpectedly observed bound to the MTase site of NSP14 (PDB: 9NJG, **Figure 3A**).

**Figure 3.**
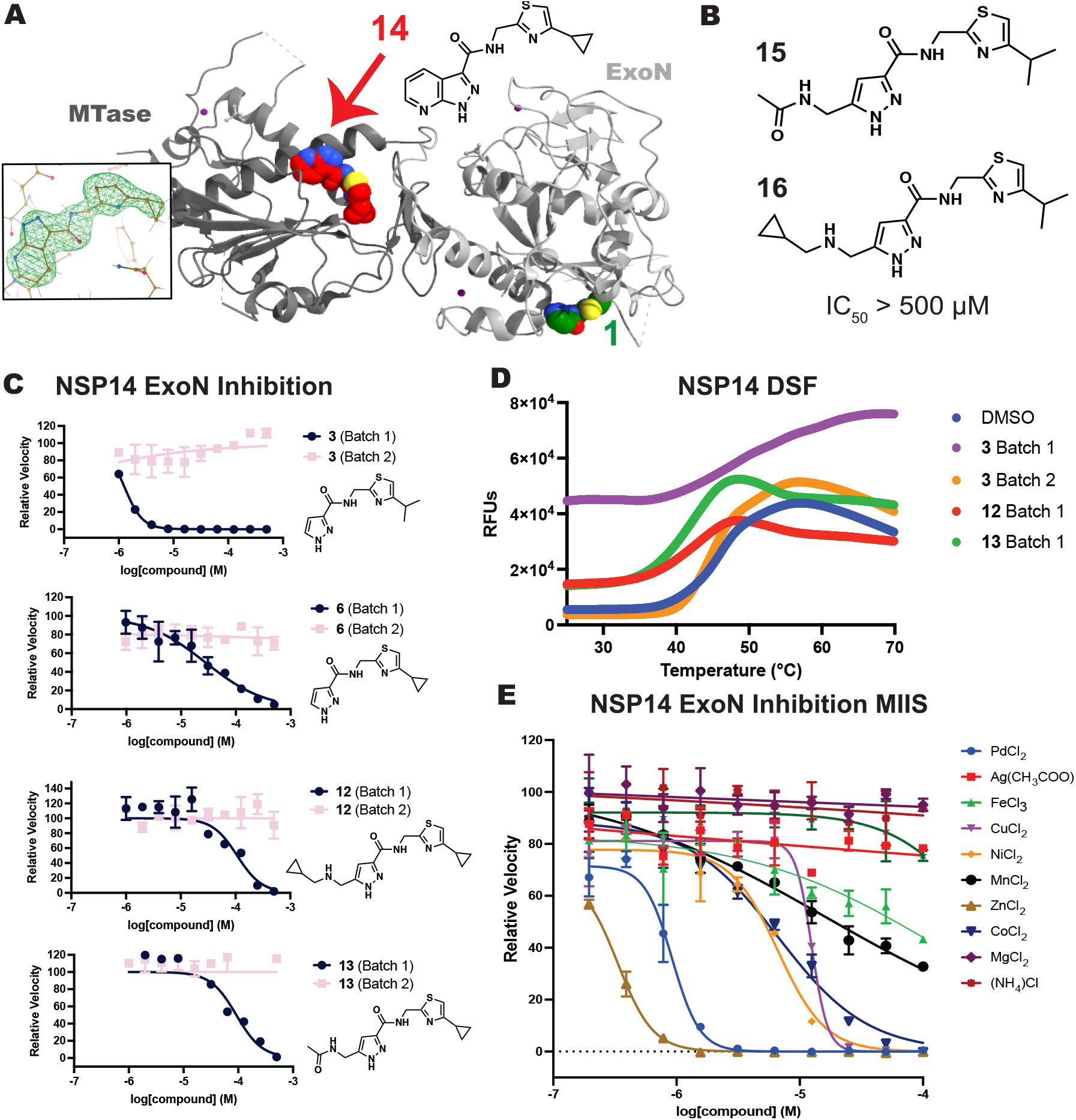
**A**. Crystal structure of **14** (red) bound to the MTase site of NSP14 (PDB: 9NJG). **B**. Inactive analogs closely related to **3. C**. Activity of two batches of **3, 6, 12**, and **13 D**. Melting curves determined for NSP14 in the presence of DMSO or 1 mM compounds. **E**. MIIS against NSP14 ExoN.

Therefore, we resynthesized larger and more stringently purified batches of **3, 6, 12**, and **13** (Supplementary Information). All second batches were inactive against NSP14 ExoN (**Figure 3C**). This null result was strengthened by our observation that the biochemically active batches of **3, 12**, and **13** displayed evidence of NSP14 destabilization by differential scanning fluorimetry (DSF), while the biochemically inactive second batch of **3** did not (**Figure 3D**).

Recently, Gerstberger *et al*. reported that metal-ion contaminants were responsible for spurious KRAS “inhibition”, even within a chemical series validated by crystallography.^25^ Because NSP14 requires two catalytic Mg^2+^ and three structural Zn^2+^ ions, we tested Gerstberger *et al*.’s Metal Ion Interference Set (MIIS) against our biochemical assay. We found that NSP14 ExoN activity is inhibited by most divalent cations (**Figure 3E**). Zn^2+^, Pd^2+^, Co^2+^, Ni^2+^, Cu^2+^, and Mn^2+^ potently inhibited NSP14 ExoN activity, with IC_50_’s equal to or more potent than our 5-substituted pyrazoles (IC_50_ 0.3 to 16 μM). Therefore, we hypothesize that contaminating divalent cations could explain the spurious biochemical activity of our original compound batches.

To our knowledge, this work represents the most substantial effort to advance the NSP14-bound fragments identified by XChem crystallographic fragment screening. We did not identify a merged compound that reproducibly inhibited NSP14 ExoN despite testing a total of eighty analogs. Before launching a hit-to-lead campaign against NSP14, practitioners should stringently remove trace metal contaminants from synthesized compounds,^25^ test multiple batches of putative inhibitors, and generate biophysical evidence of binding in solution. Given the promising genetic validation of NSP14 ExoN as an antiviral target, we hope that this work will inspire future efforts to inhibit NSP14 ExoN via an alternative strategy.

## MATERIALS AND METHODS

### Purification and Crystallization of NSP14

Near full-length SARS-CoV-2 NSP14 (Leu5932—Gln6452; UniProt P0DTD1) with a TEV-cleavable, N-terminal His6-ZB tag was obtained as a gift in the pNIC-ZB backbone from Imprachim, Yosaatmadja, and Newman and was purified and crystallized essentially as previously reported.^5^ The construct was transformed into Rosetta 2 (DE3) competent cells (Sigma-Aldrich #71397) and expressed in TB-PLUS media supplemented with 0.5 mM ZnCl_2_.^26^ Cells were lysed by sonication in Lysis Buffer (Base Buffer + 1X Roche cOmplete™ Protease Inhibitors + 0.1 mg/mL egg-white lysozyme + 1X benzonase) and purified by IMAC on HisPur NiNTA resin (Thermo Fisher). The His6-ZB tag was cleaved during overnight dialysis into Gel Filtration Buffer with TEV protease (1:10 w:w) at 4°C and removed via reverse IMAC on NiNTA resin. Cleaved NSP14 was polished on a Superdex 200 column (GE) into Gel Filtration Buffer and Monodisperse NSP14 was concentrated to 10 mg/mL prior to long-term storage at −80°C. <u>Base Buffer:</u> 50 mM HEPES pH = 7.5, 500 mM NaCl, 5% glycerol, 1 mM TCEP. <u>Gel Filtration Buffer:</u> 20 mM HEPES pH = 7.0, 500 mM NaCl, 5% glycerol, 0.5 mM TCEP.

Sitting drop crystallization plates were set at 1:1, 1:2, and 2:1 v:v (protein:reservoir) ratios in 150 nL drops by a Mosquito Xtal 3 (SPT Labtech) using a custom screen. Crystals, which typically appeared after 14 days in the 1:1 drops between 0.18—0.22M K_2_HPO_4_ and 1.4—1.6M NaH_2_PO4, were soaked with ligands for 2 hours (final concentration 10— 25 mM, 10% DMSO) prior to freezing without additional cryoprotection in LN_2_. Automated diffraction data was collected at Diamond Lightsource Beamline I03 and I04. Structures were indexed and merged using Xia2/DIALS, solved by molecular replacement with Phaser using PDB: 5SKY (with the ligand **1** removed) as a search model, refined with Phenix, and manually rebuilt in Coot where necessary.^27-32^ Figures were rendered in ChimeraX.^33^ Data collection and refinement statistics are provided in the **Supplemental Information**, and all structures have been deposited to the PDB: 9NAZ, 9NFP, 9NHA, 9NIO, 9NJG, 9NHU.

### Purification of NSP10

Full-length SARS-CoV-2 NSP10 (Ala4254—Gln4932; UniProt P0DTD1) with a 3C-cleavable, N-terminal GST tag was obtained as a gift in the pNIC-CTH0 backbone from Imprachim, Yosaatmadja, and Newman and purified essentially as previously reported.^5^ Construct was expressed in TB-PLUS and lysed as described above. GST-NSP10 was purified over Glutathione® Sepharose (Sigma) in Base Buffer and then the tag was removed with His6-HRV3C protease (1:20 w:w) during overnight dialysis into Gel Filtration Buffer (20 mM HEPES pH = 8.0, 500 mM NaCl, 5% glycerol, 0.5 mM TCEP). Cleaved protein was passed over a Talon® column (Takara) to remove the His6-HRV3C and NSP10 was polished on a Superdex 75 (GE) equilibrated in Gel Filtration Buffer. Monodisperse peak was pooled and stored at −80°C at 2-3 mg/mL.

### RNA Hydrolysis TR-FRET ExoN Activity Assay

Two RNA oligos, the 5’ FAM conjugated top strand (5’-FAM-ACUAAUAAUAUCA) and the 3’ BHQ-1 conjugated bottom strand (AAAUAGAUAUUAUUAGU-3’BHQ-1) (IDT) were annealed in annealing buffer (100 mM NaCl, 20 mM HEPES-NaOH pH 7.5, 0.2 mM MgCl_2_) at 12 µM by heating to 95 °C for 5 min in aC1000 Touch™ thermal cycler and then cooling to 26 °C. 30 µL reactions in ExoN Assay Buffer (100 mM NaCl, 20 mM HEPES-NaOH pH = 7.5, 2 mM MgCl_2_, 0.01% TritonX-100, 1 mM TCEP) were conducted black non-binding 384-well plates (Corning 3575). Compound dilutions, prepared in 100% DMSO, were stamped into each well with an ECHO^®^ 550 liquid handler and then 20 µL NSP14 (83.3 nM final) was added and incubated for 15 min at room temperature. 5 µl of NSP10 (83.3 nM final, 1:1 ratio) was added and incubated at 37^°^C for 20 min. Finally, 5 µL dsRNA substrate 5 µL was added (208.3 nM final) and a kinetic time-course for FAM flouresence was collected on a BioTek Cytation 5 at 37^°^C. Initial rates were extracted by linear regression and then normalized to control wells with DMSO (100% activity) or no NSP10 (0% activity) prior to IC_50_ determination by non-linear regression in GraphPad PRISM.

### Counter-Screening Assays

RNAse T1 inhibition assay and thiazole orange RNA intercalation counter-screening assays were conducted essentially as previously reported.^24^ The TR-FRET ExoN activity assay dsRNA substrate and Assay Buffer were used for the intercalation assay and mixed with compounds at a final 0.5% DMSO.

### NSP14 DSF Thermal Shift Assay

In technical triplicate, NSP14 (0.15 mg/mL) was prepared in 20 mM HEPES pH = 7.5, 200 mM NaCl, 1 mM TCEP, 2 mM MgCl_2_ and mixed with 1 mM compound or 2% DMSO. Samples were rested for 10 min and then mixed with SYPRO™ Orange (3X concentration, Invitrogen #S6650). Raw melting curves (25-99 °C, 0.05 °C C/sec, 20 uL volume) were collected on a QuantStudio 3 (Applied Biosystems) and each replicate well was individually normalized from 0-100 prior to averaging.

### Compound Synthesis and Characterization

See supplemental information.

## ACKNOWLEDGEMENTS

This research was supported by an agreement between the Advanced Photon Source, a U.S. Department of Energy (DOE) Office of Science user facility operated for the DOE Office of Science by Argonne National Laboratory under Contract No. DE-AC02-06CH11357, and the Diamond Light Source, the U.K.’s national synchrotron science facility, located at the Harwell Science and Innovation Campus in Oxfordshire, where the work was performed under proposal AU34221.

