## Supplemental Information for "Fragment-Based Development of NSP14 Exonuclease Inhibitors Confounded by Batch-to-Batch Variability"

**Crystallography Data Collection and Refinement Statistics (Table 1)**

| | **PDB: 9NIO (Compound 4)** |
|---|---|
| **Wavelength** | |
| **Resolution range** | 48.58 - 2.0 (2.072 - 2.0) |
| **Space group** | P 21 21 21 |
| **Unit cell** | 67.5846 68.3691 138.088 90 90 90 |
| **Total reflections** | 600438 (61052) |
| **Unique reflections** | 44014 (4316) |
| **Multiplicity** | 13.6 (14.1) |
| **Completeness (%)** | 99.88 (99.81) |
| **Mean I/sigma(I)** | 11.05 (1.32) |

| | |
|---|---|
| **Wilson B-factor** | 31.93 |
| **R-merge** | 0.1558 (1.371) |
| **R-meas** | 0.1619 (1.422) |
| **R-pim** | 0.04381 (0.377) |
| **CC1/2** | 0.995 (0.868) |
| **CC*** | 0.999 (0.964) |
| **Reflections used in refinement** | 43966 (4308) |
| **Reflections used for R-free** | 2008 (199) |
| **R-work** | 0.2084 (0.2656) |
| **R-free** | 0.2299 (0.2925) |
| **CC(work)** | 0.934 (0.906) |
| **CC(free)** | 0.928 (0.850) |
| **Number of non-hydrogen atoms** | 3625 |
| **macromolecules** | 3394 |
| **ligands** | 37 |
| **solvent** | 202 |
| **Protein residues** | 426 |
| **RMS(bonds)** | 0.010 |
| **RMS(angles)** | 1.03 |

| | |
|---|---|
| **Ramachandran favored (%)** | 97.57 |
| **Ramachandran allowed (%)** | 2.43 |
| **Ramachandran outliers (%)** | 0.00 |
| **Rotamer outliers (%)** | 0.00 |
| **Clashscore** | 2.69 |
| **Average B-factor** | 36.41 |
| **macromolecules** | 35.99 |
| **ligands** | 52.53 |
| **solvent** | 41.17 |

| | **PDB: 9NAZ (Compound 5)** |
|---|---|
| **Wavelength** | |
| **Resolution range** | 60.84 - 2.3 (2.382 - 2.3) |
| **Space group** | P 21 21 21 |
| **Unit cell** | 67.7547 68.7281 138.245 90 90 90 |
| **Total reflections** | 345732 (20223) |
| **Unique reflections** | 29271 (2808) |
| **Multiplicity** | 11.8 (7.2) |
| **Completeness (%)** | 99.45 (97.67) |
| **Mean I/sigma(I)** | 10.19 (1.16) |

| | |
|---|---|
| **Wilson B-factor** | 35.11 |
| **R-merge** | 0.1449 (1.297) |
| **R-meas** | 0.1512 (1.391) |
| **R-pim** | 0.04248 (0.4854) |
| **CC1/2** | 0.996 (0.816) |
| **CC*** | 0.999 (0.948) |
| **Reflections used in refinement** | 29237 (2803) |
| **Reflections used for R-free** | 2003 (192) |
| **R-work** | 0.2111 (0.2652) |
| **R-free** | 0.2397 (0.2976) |
| **CC(work)** | 0.922 (0.856) |
| **CC(free)** | 0.880 (0.893) |
| **Number of non-hydrogen atoms** | 3679 |
| **macromolecules** | 3473 |
| **ligands** | 39 |
| **solvent** | 177 |
| **Protein residues** | 433 |
| **RMS(bonds)** | 0.014 |
| **RMS(angles)** | 1.28 |

| | |
|---|---|
| **Ramachandran favored (%)** | 97.86 |
| **Ramachandran allowed (%)** | 2.14 |
| **Ramachandran outliers (%)** | 0.00 |
| **Rotamer outliers (%)** | 1.06 |
| **Clashscore** | 2.48 |
| **Average B-factor** | 38.68 |
| **macromolecules** | 38.42 |
| **ligands** | 47.41 |
| **solvent** | 42.28 |

16

| | **PDB: 9NFP (Compound 6)** |
|---|---|
| **Wavelength** | |
| **Resolution range** | 60.81 - 2.3 (2.382 - 2.3) |
| **Space group** | P 21 21 21 |
| **Unit cell** | 67.7267 68.4265 138.082 90 90 90 |
| **Total reflections** | 320692 (31988) |
| **Unique reflections** | 28946 (2856) |
| **Multiplicity** | 11.1 (11.2) |
| **Completeness (%)** | 98.88 (99.89) |
| **Mean I/sigma(I)** | 20.36 (7.39) |

| | |
|---|---|
| **Wilson B-factor** | 32.35 |
| **R-merge** | 0.0839 (0.3837) |
| **R-meas** | 0.08802 (0.402) |
| **R-pim** | 0.02615 (0.118) |
| **CC1/2** | 0.998 (0.981) |
| **CC*** | 0.999 (0.995) |
| **Reflections used in refinement** | 28919 (2853) |
| **Reflections used for R-free** | 1447 (133) |
| **R-work** | 0.2060 (0.2212) |
| **R-free** | 0.2305 (0.2639) |
| **CC(work)** | 0.932 (0.947) |
| **CC(free)** | 0.931 (0.894) |
| **Number of non-hydrogen atoms** | 3614 |
| **macromolecules** | 3358 |
| **ligands** | 42 |
| **solvent** | 226 |
| **Protein residues** | 420 |
| **RMS(bonds)** | 0.005 |
| **RMS(angles)** | 0.73 |

| | |
|---|---|
| **Ramachandran favored (%)** | 97.55 |
| **Ramachandran allowed (%)** | 2.45 |
| **Ramachandran outliers (%)** | 0.00 |
| **Rotamer outliers (%)** | 0.27 |
| **Clashscore** | 1.36 |
| **Average B-factor** | 36.87 |
| **macromolecules** | 36.34 |
| **ligands** | 45.41 |
| **solvent** | 43.55 |

17

| | **PDB: 9NHU (Compound 12)** |
|---|---|
| **Wavelength** | |
| **Resolution range** | 60.83 - 2.1 (2.175 - 2.1) |
| **Space group** | P 21 21 21 |
| **Unit cell** | 67.7903 68.2164 137.873 90 90 90 |
| **Total reflections** | 517456 (52509) |
| **Unique reflections** | 38102 (3739) |
| **Multiplicity** | 13.6 (14.0) |
| **Completeness (%)** | 97.93 (87.98) |
| **Mean I/sigma(I)** | 5.70 (0.41) |

| | |
|---|---|
| **Wilson B-factor** | 39.66 |
| **R-merge** | 0.3504 (4.643) |
| **R-meas** | 0.3641 (4.818) |
| **R-pim** | 0.0982 (1.278) |
| **CC1/2** | 0.984 (0.361) |
| **CC*** | 0.996 (0.728) |
| **Reflections used in refinement** | 37322 (3294) |
| **Reflections used for R-free** | 1837 (175) |
| **R-work** | 0.2317 (0.3779) |
| **R-free** | 0.2721 (0.3996) |
| **CC(work)** | 0.940 (0.674) |
| **CC(free)** | 0.911 (0.709) |
| **Number of non-hydrogen atoms** | 3518 |
| **macromolecules** | 3370 |
| **ligands** | 57 |
| **solvent** | 112 |
| **Protein residues** | 423 |
| **RMS(bonds)** | 0.009 |
| **RMS(angles)** | 0.97 |

| | |
|---|---|
| **Ramachandran favored (%)** | 96.84 |
| **Ramachandran allowed (%)** | 2.92 |
| **Ramachandran outliers (%)** | 0.24 |
| **Rotamer outliers (%)** | 0.00 |
| **Clashscore** | 3.16 |
| **Average B-factor** | 45.40 |
| **macromolecules** | 45.26 |
| **ligands** | 59.31 |
| **solvent** | 44.97 |

18

| | **PDB: 9NJG (Compound 14)** |
|---|---|
| **Wavelength** | |
| **Resolution range** | 60.81 - 2.1 (2.175 - 2.1) |
| **Space group** | P 21 21 21 |
| **Unit cell** | 67.7308 67.654 138.695 90 90 90 |
| **Total reflections** | 513354 (51861) |
| **Unique reflections** | 37986 (3758) |
| **Multiplicity** | 13.5 (13.8) |
| **Completeness (%)** | 99.73 (99.71) |
| **Mean I/sigma(I)** | 9.52 (0.85) |

| | |
|---|---|
| **Wilson B-factor** | 37.96 |
| **R-merge** | 0.1604 (1.725) |
| **R-meas** | 0.1668 (1.791) |
| **R-pim** | 0.04526 (0.4797) |
| **CC1/2** | 0.998 (0.835) |
| **CC*** | 1 (0.954) |
| **Reflections used in refinement** | 37887 (3747) |
| **Reflections used for R-free** | 2003 (196) |
| **R-work** | 0.2362 (0.3590) |
| **R-free** | 0.2741 (0.4121) |
| **CC(work)** | 0.926 (0.867) |
| **CC(free)** | 0.902 (0.714) |
| **Number of non-hydrogen atoms** | 3321 |
| **macromolecules** | 3172 |
| **ligands** | 47 |
| **solvent** | 115 |
| **Protein residues** | 401 |
| **RMS(bonds)** | 0.012 |
| **RMS(angles)** | 1.17 |

| | |
|---|---|
| **Ramachandran favored (%)** | 97.40 |
| **Ramachandran allowed (%)** | 2.60 |
| **Ramachandran outliers (%)** | 0.00 |
| **Rotamer outliers (%)** | 0.29 |
| **Clashscore** | 2.88 |
| **Average B-factor** | 44.89 |
| **macromolecules** | 44.68 |
| **ligands** | 51.79 |
| **solvent** | 48.51 |

*Statistics for the highest-resolution shell are shown in parentheses.*

### Compound Synthesis and Characterization

**General Chemistry**. All chemical reagents and reaction solvents were purchased from commercial suppliers and used as received. Normal phase chromatography was performed on a Teledyne ISCO CombiFlash NextGen300 system using Teledyne RediSep® normal phase silica cartridges, with average particle size 35-70 micron. Preparative reversed-phase HPLC was performed using either Gilson GX-271 preparative liquid handler, or Teledyne ACCQ-Prep purification system. Both instruments were equipped with Phenomenex Kinetex or Gemini 5 micron C18 100 x 33 mm columns, using gradients of MeCN in H2O as mobile phase with 0.1% TFA modifier. Flow rate 40 mL/min, gradient 5–40% organic phase B over a 14 min gradient, then 100% B for 3 min. A: $H_2O$ + 0.1% TFA; B: MeCN + 0.1% TFA. Compounds that are obtained as a TFA salt after purification were afforded as free base, by dissolving the salt in EtOAc and washing with sat. aq. $K_2CO_3$, or by elution through a Biotage ISOLUTE SCX-II cartridge, loading and washing with MeOH and eluting with 2N $NH_3$ in MeOH. Proton nuclear magnetic resonance ($^1$H NMR) spectra were recorded at 400 MHz with standard pulse sequences. For $^1$H NMR spectra, chemical shifts are reported in parts per million (ppm) and are reported relative to residual non-deuterated solvent signals. Coupling constants are reported in Hertz (Hz). The following abbreviations (or a combination, thereof) are used to describe splitting patterns: s, singlet; d, doublet; t, triplet; q, quartet; pent, pentet; m, multiplet; br, broad. Analytical thin layer chromatography (TLC) was performed on Kieselgel 60 F254 glass plates precoated with a 0.25 mm thickness of silica gel. TLC plates were visualized with UV light and iodine. Hydrogenation reactions are performed using an atmospheric balloon. All compounds were determined to be at 95% purity or higher, unless otherwise noted, as measured by analytical reversed-phase HPLC with either an Agilent or Shimadzu RP-HPLC LCMS system. Detection methods are diode array (DAD) at 210, 254 nM and positive/negative electrospray ionization (ESI), mass range capable of 25-20,000 *m/z*. The MS detectors are configured to mass range 100 to 1700 *m/z*, with nitrogen used as nebulizer gas. Unless stated, all methods use an Agilent InfinityLab Poroshell 120 EC-C18 column, dimensions 4.6 x 50 mm, 2.7 μm, fitted with

Poroshell 120 EC-C18, 2.1 mm, 1.9 µm guard. Mobile phase A was 0.1% TFA in $H_2O$, mobile phase B was 0.1% TFA in $CH_3CN$.

The following compounds were purchased from Enamine and used without further purification, purity per supplier >95%: 2,4-dimethyl-6-(piperazin-1-yl)pyrimidine catalog #Z274575916 (**1**), *N*-(thiazol-2-ylmethyl)-7*H*-pyrrolo[2,3-*d*]pyrimidine-5-carboxamide catalog #Z5068737199 (**8**), *N*-(2-(4,5-dimethylthiazol-2-yl)propan-2-yl)-7*H*-pyrrolo[2,3-*d*]pyrimidine-5-carboxamide catalog #Z3297671643 (**9**), *N*-((4-cyclopropylthiazol-2-yl)methyl)-5-(2,4-dimethylphenyl)-1*H*-pyrazole-3-carboxamide catalog #Z1747165378 (**10**), *N*-((4-cyclopropylthiazol-2-yl)methyl)-5-(pyridin-4-yl)-1*H*-pyrazole-3-carboxamide catalog #Z3540570837 (**11**).

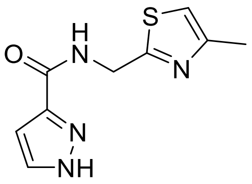

***N*-((4-Methylthiazol-2-yl)methyl)-1*H*-pyrazole-3-carboxamide (2).**

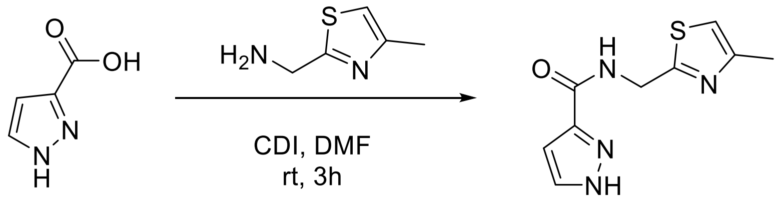


**2**

**Scheme 1.** Synthetic route to prepare **2**.

To a solution of commercially available 1*H*-pyrazole-3-carboxylic acid (30.0 mg, 0.3 mmol, CAS 1621-91-6) in DMF (3.0 mL), under $N_2$, were added 1,1'-carbonyldiimidazole (CDI, 87.6 mg, 0.5 mmol). The reaction mixture was then stirred at rt under $N_2$ for 0.5 hours. 4-Methylthiazol-2-yl)methanamine (34.2 mg, 0.3 mmol, CAS 144163-68-8) was then added. Upon completion of the reaction as evident by LCMS the mixture was purified by RP-HPLC to afford *N*-((4-methylthiazol-2-yl)methyl)-1*H*-pyrazole-3-carboxamide (35 mg,

59%) as a white solid: LCMS [M+H-$CF_3COOH$] 223.0; HPLC ≥95% (254 nm); $^1$H NMR (400 MHz, DMSO) δ 13.30 (s, 1H), 8.94 (s, 1H), 7.84 (s, 1H), 7.11 (s, 1H), 6.67 (s, 1H), 4.63 (d, J = 6.2 Hz, 2H), 2.32 (s, 3H).

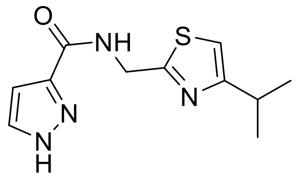

***N*-((4-Isopropylthiazol-2-yl)methyl)-1*H*-pyrazole-3-carboxamide (3)**.

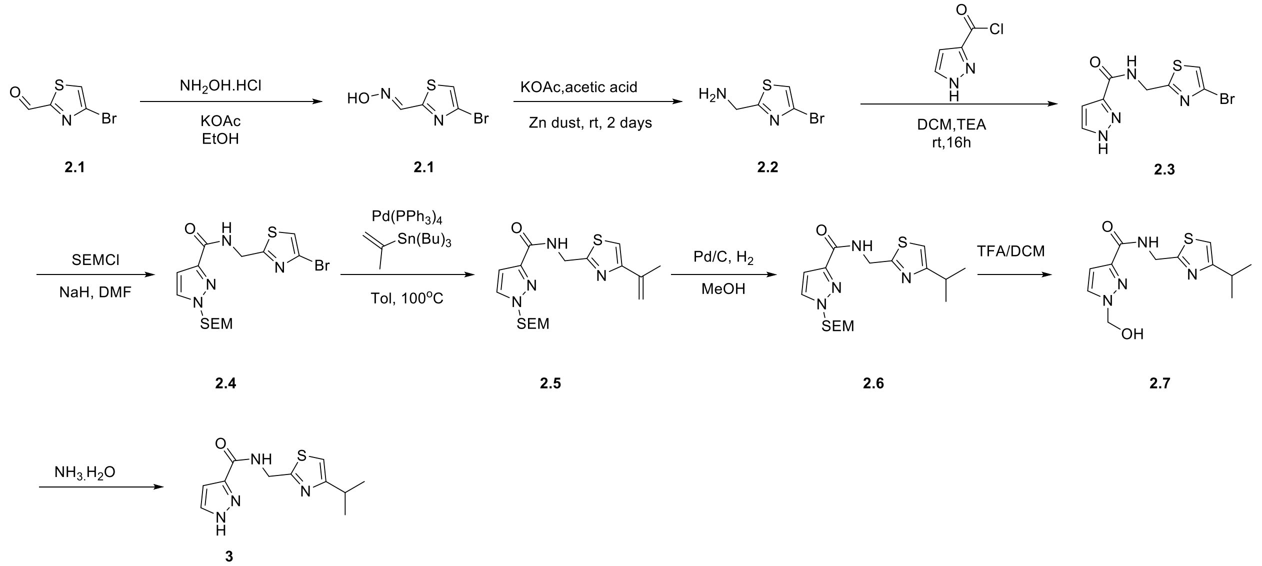

**Scheme 2.** Batch 1 synthetic route compound **3.**

*Step 1. 4-Bromothiazole-2-carbaldehyde oxime (**2.1**).*

To a solution of 4-bromothiazole-2-carboxaldehyde (5.0 g, 26.0 mmol) in 30 mL EtOH, was added $NH_2OH$ (10 mL). The mixture was stirred at 60°C for 3h under $N_2$. The mixture was concentrated to afford a crude product. The mixture was extracted with EtOAc, washed by 0.2 M HCl. Compound **2.1** was obtained as a yellow solid (5.2 g, 99%): LCMS [M+H] 209.2.

*Step 2. (4-Bromothiazol-2-yl)methanamine (**2.2**).*

To a solution of 4-bromothiazole-2-carbaldehyde oxime (2.6 g, 12.5 mmol) in 50 mL acetic acid were added KOAc (2.5 g, 25 mmol) and Zn dust (2.4 g, 37 mmol) at rt under $N_2$ for overnight. The mixture was filtered. After washing with $NaHCO_3$, the organics were extracted with DCM/MeOH (10:1), dried over $Na_2SO_4$, and concentrated to dryness to give **2.2** as a yellow solid (1.5 g, 63%): LCMS [M+H] 193.2.

*Step 3. N-((4-Bromothiazol-2-yl)methyl)-1H-pyrazole-3-carboxamide (**2.3**).*

To a solution of (4-bromothiazol-2-yl)methanamine (780 mg, 4.1 mmol) in 10 mL DCM was added TEA (1.2 g, 12.2 mmol) at rt. The mixture was cooled to 0$^o$C and 1*H*-pyrazole-3-carbonyl chloride was (528 mg, 4.1 mmol) added slowly. The mixture was stirred at 25$^o$C for 4h under $N_2$. After washing with $H_2O$, the organics were extracted with DCM, dried over $Na_2SO_4$, and concentrated to dryness. The crude product was purified by automated silica gel chromatography DCM : MeOH (40:1) and dried to give **2.3** as a yellow solid (570 mg, 49%): LCMS [M+H] 288.9, ≥95% (214, 254 nm).

*Step 4. N-((4-Bromothiazol-2-yl)methyl)-1-((2-(trimethylsilyl)ethoxy)methyl)-1H-pyrazole-3-carboxamide (**2.4**).*

To a solution of *N*-((4-bromothiazol-2-yl)methyl)-1*H*-pyrazole-3-carboxamide (2.2 g, 7.7 mmol, 1.0 eq) in 20 mL DMF was added NaH (185 mg, 7.7 mmol, 1.0 eq) at 0$^o$C under $N_2$ and stirred for 0.5 h. SEMCl (1.3 g, 7.7 mmol, 1.0 eq) was added. The mixture was left at 0 $^o$C for 4 h under $N_2$. The mixture was extracted with EtOAc and the organic layers dried over $Na_2SO_4$. The crude product was purified by automated silica gel chromatography to give intermediate **2.4** as a yellow solid (3.0 g, 91%): LCMS [M+H] 419.1, ≥95% (214, 254 nm).

*Step 5. N-((4-(Prop-1-en-2-yl)thiazol-2-yl)methyl)-1-((2-(trimethylsilyl)ethoxy)methyl)-1H-pyrazole-3-carboxamide (**2.5**).*

To a solution of *N*-((4-bromothiazol-2-yl)methyl)-1-((2-(trimethylsilyl)ethoxy)methyl)-1*H*-pyrazole-3-carboxamide (500 mg, 1.2 mmol) in 5 mL dioxane were added $Pd(Ph_3)_4$ (69 mg, 0.06 mmol), tributyl(prop-1-en-2-yl)stannane (1.2 g, 3.6 mmol). The mixture was stirred at 110$^o$C for 16 h under $N_2$. The mixture was then extracted with EtOAc (3x), the organic layers combined and dried over $Na_2SO_4$ and concentrated to dryness. The crude product was purified by RP-HPLC to give a white solid (82 mg, 18%): LCMS [M+H] 379.2, ≥95% (214, 254 nm).

*Step 6. N-((4-Isopropylthiazol-2-yl)methyl)-1-((2-(trimethylsilyl)ethoxy)methyl)-1H-pyrazole-3-carboxamide (**2.6**).*

To a solution of *N*-((4-(prop-1-en-2-yl)thiazol-2-yl)methyl)-1-((2-(trimethylsilyl)ethoxy)methyl)-1*H*-pyrazole-3-carboxamide (82 mg, 0.2 mmol) in 2 mL MeOH were added Pd/C (8 mg), $Pd(OH)_2$ (8 mg), and HCl-MeOH (2 drops) at rt. The mixture was stirred for 3h under an atmospheric $H_2$ balloon, filtered over Celite, and the filtrate concentrated. The crude product **2.6** was used directly in the next step without further purification: LCMS [M+H] 381.1, ≥95% (214, 254 nm).

*Step 7. 1-(Hydroxymethyl)-N-((4-isopropylthiazol-2-yl)methyl)-1H-pyrazole-3-carboxamide (**2.7**).*

To a solution of *N*-((4-isopropylthiazol-2-yl)methyl)-1-((2-(trimethylsilyl)ethoxy)methyl)-1*H*-pyrazole-3-carboxamide (80 mg, 0.2 mmol) in 2 mL DCM were added TFA (0.5 mL) for 2h. The mixture was extracted with DCM (2x), the organic phases dried over $Na_2SO_4$ and concentrated to dryness. The crude product **2.7** was used directly in the next step without further purification: LCMS [M+H] 281.1, ≥95% (214, 254 nm).

*Step 8. N-((4-Isopropylthiazol-2-yl)methyl)-1H-pyrazole-3-carboxamide (**3**).*

To a solution of 1-(hydroxymethyl)-*N*-((4-isopropylthiazol-2-yl)methyl)-1H-pyrazole-3-carboxamide (**2.7**, 59 mg, 0.2 mmol, 1.0 eq) in 2 mL MeOH were added $NH_3 \cdot H_2O$ (0.2 mL) and $Pd(PPh_3)_4$ at rt for 1h. The mixture was purified by RP-HPLC to give title compound as a white solid (15.8 mg, 30%): LCMS [M+H] 251.1; HPLC ≥95% (254 nm); $^1$H NMR (400 MHz, DMSO) δ 9.03 (s, 1H), 7.80 (d, J=2.4 Hz, 1H), 7.11 (s, 1H), 6.71 (d, J=2.4 Hz, 1H), 4.65 (d, J=6.0 Hz, 2H), 3.00-2.97 (m, 1H), 1.23 (d, J=6.8 Hz, 6H).

***Batch 2 Synthesis Compound 3.***

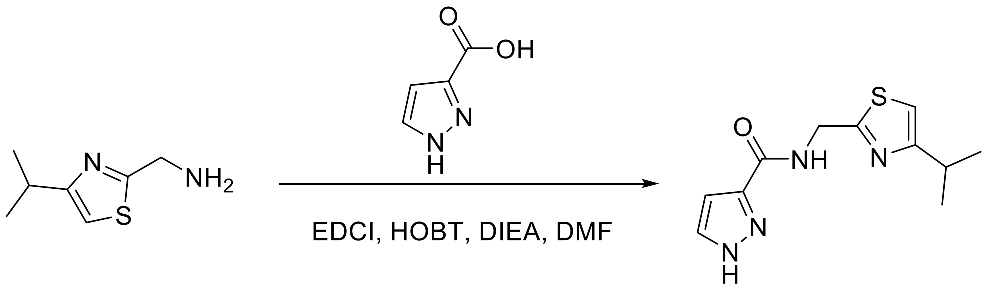


**3** (batch 2)

***Preparation of N-((4-isopropylthiazol-2-yl)methyl)-1H-pyrazole-3-carboxamide using commercially available (4-isopropylthiazol-2-yl)methanamine (3).***

To a solution of commercially available (4-isopropylthiazol-2-yl)methanamine (70 mg, 0.448 mmol, CAS 643725-72-8) 1*H*-pyrazole-3-carboxylic acid (50 mg, 0.448 mmol, CAS 1621-91-6), *N*,*N*-diisopropylethylamine (115 mg, 0.897 mmol) in DMF was added EDCI (103 mg, 0.538 mmol) and HOBt (72 mg, 0.538 mmol). The mixture was stirred at 50°C for 1 h under $N_2$. The progress of the reaction was monitored by LCMS. After complete conversion of starting material, 10 mL of EtOAc was added, the organic solvent washed with $H_2O$ (10 mL), followed by brine (20 mL). The organic phase was dried over $Na_2SO_4$ and concentrated under reduced pressure. The crude product was purified by RP-HPLC to give *N*-((4-isopropylthiazol-2-yl)methyl)-1*H*-pyrazole-3-carboxamide (**3**) as a white solid (32 mg, 29%): LCMS [M+H] 251.0; HPLC ≥95% (254 nm); $^1$H NMR (400 MHz, DMSO) δ 9.02 (s, 1H), 7.78 (t, J = 13.3 Hz, 1H), 7.11 (s, 1H), 6.71 (d, J = 2.1 Hz, 1H), 4.65 (d, J = 6.2 Hz, 2H), 3.06 – 2.93 (m, 1H), 1.23 (d, J = 6.9 Hz, 6H).

***Batch 1 NMR***

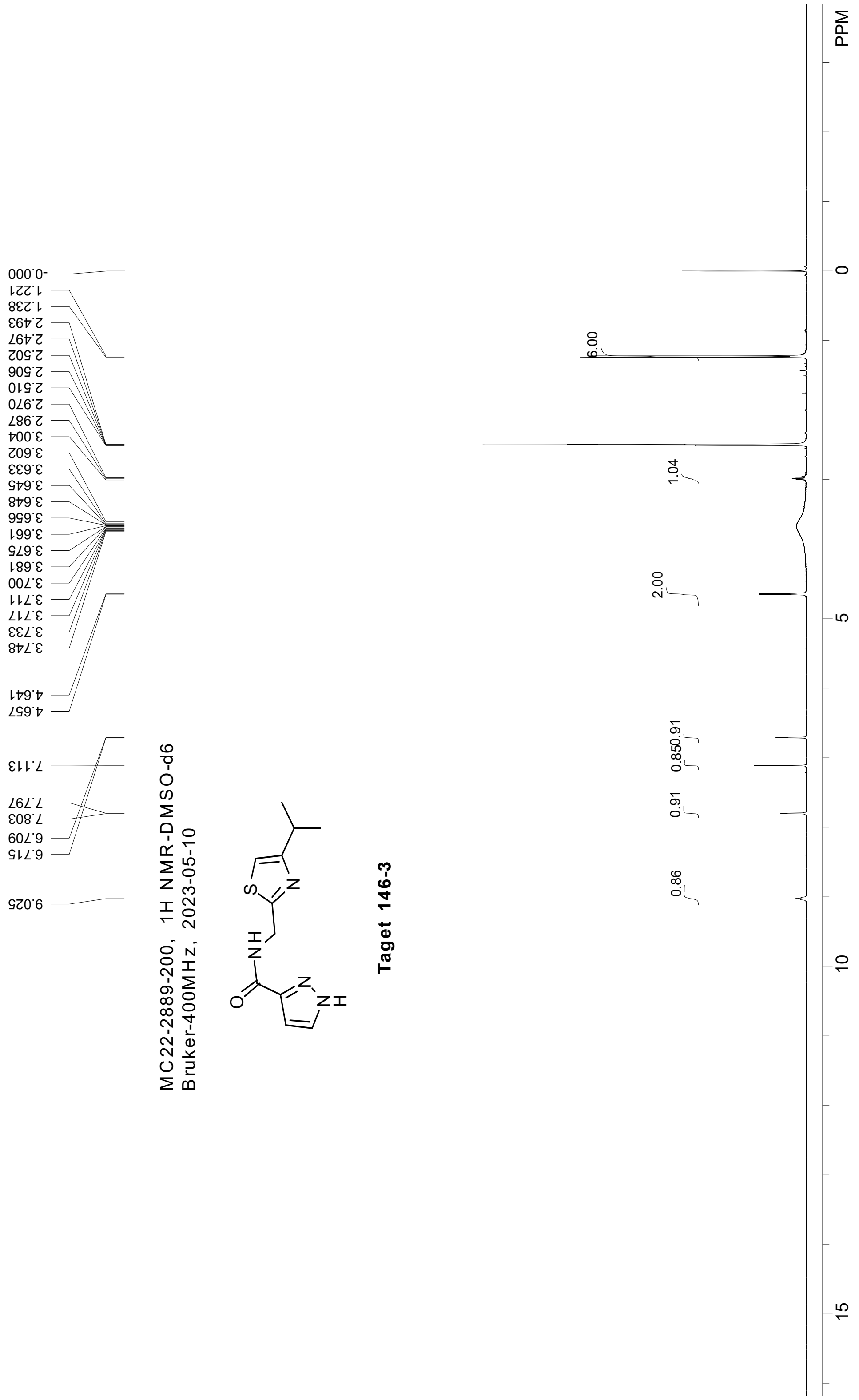
-0.000
1.221
1.238
2.493
2.497
2.502
2.506
2.510
2.970
2.987
3.004
3.602
3.633
3.645
3.648
3.656
3.661
3.675
3.681
3.700
3.711
3.717
3.733
3.748
4.641
4.657
7.113
7.797
7.803
6.709
6.715
9.025
MC22-2889-200, 1H NMR-DMSO-d6
Bruker-400MHz, 2023-05-10
Taget 146-3
6.00
1.04
2.00
0.850.91
0.91
0.86
PPM
0
5
10
15

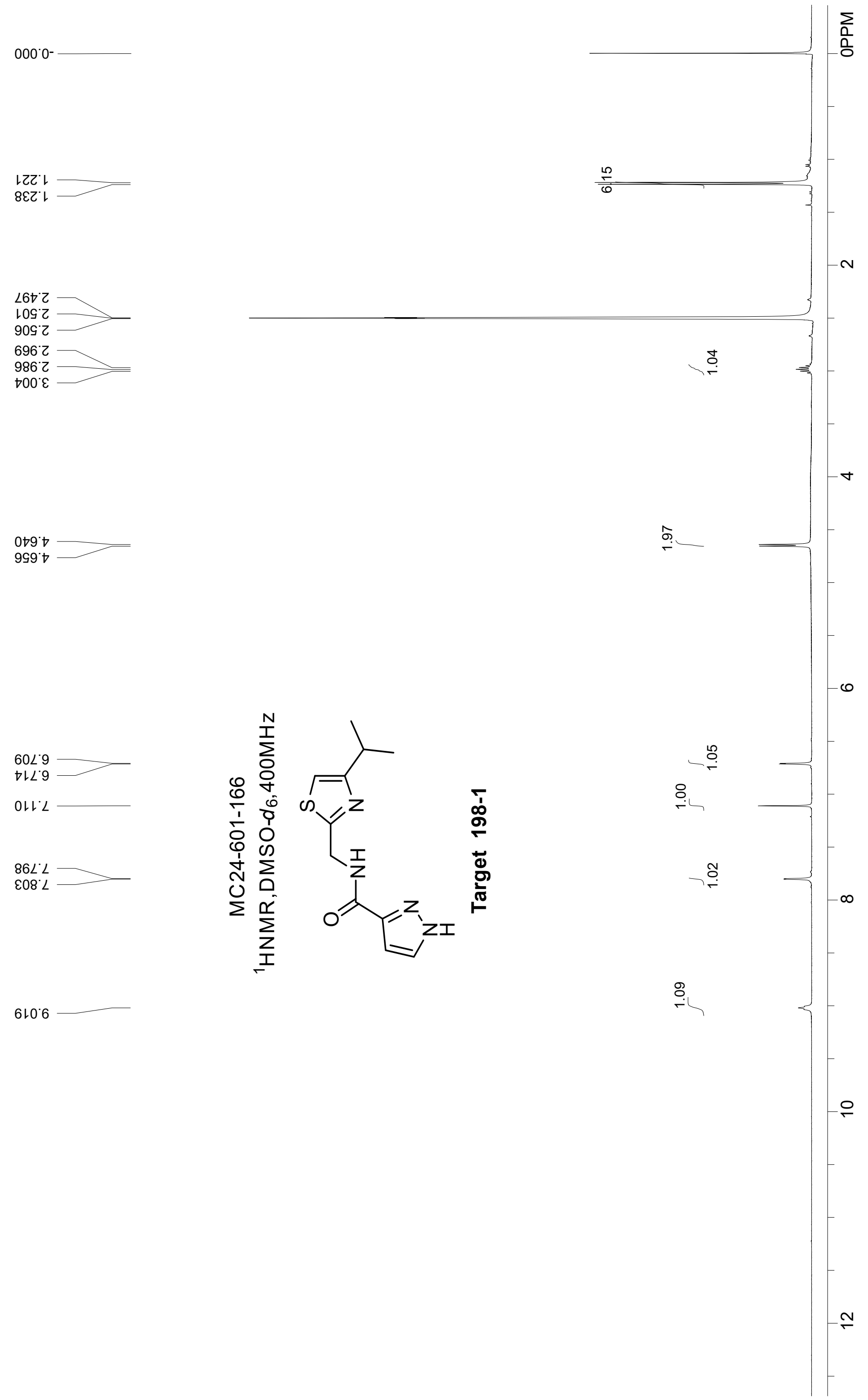
MC24-601-166
$^1$HNMR,DMSO-$d_6$,400MHz
Target 198-1
-0.000
1.221
1.238
2.497
2.501
2.506
2.969
2.986
3.004
4.640
4.656
6.709
6.714
7.110
7.798
7.803
9.019
6.15
1.04
1.97
1.05
1.00
1.02
1.09
0PPM
2
4
6
8
10
12

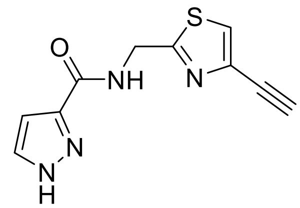

**_N_-((2-Ethynylthiazol-4-yl)methyl)-1_H_-pyrazole-3-carboxamide (4).**

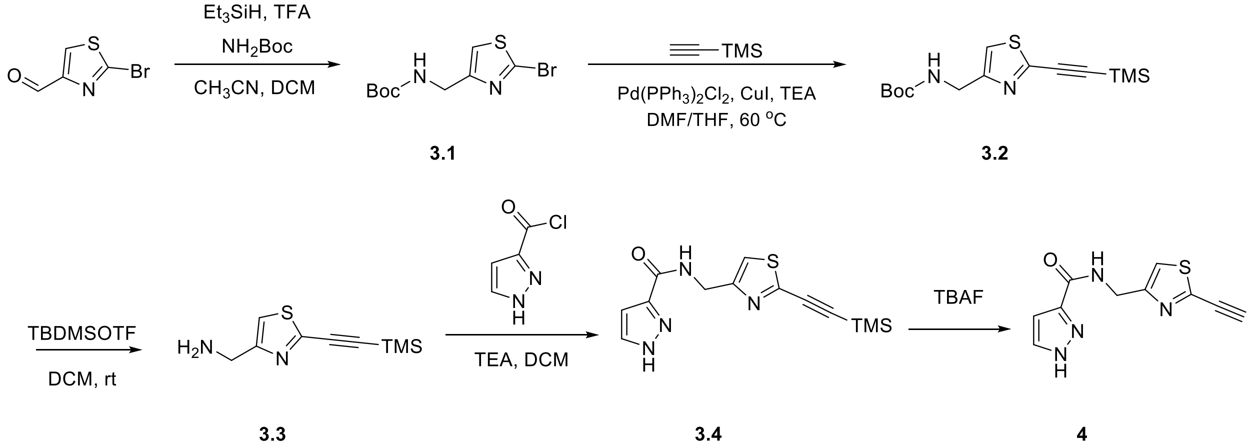


**Scheme 3.** Synthetic route compound **4**.

*Step 1. Tert-butyl ((2-bromothiazol-4-yl)methyl)carbamate (**3.1**).*

To a solution of 2-bromothiazole-4-carbaldehyde (800 mg, 4.2 mmol) in 16 mL $CH_3CN$:DCM=1:1, were added $BocNH_2$ (980 mg, 8.4 mmol), TFA (477 mg, 4.2 mmol), $Et_3SiH$ (971 mg, 8.4 mmol). The mixture was left at 60 °C for 6h under $N_2$. The mixture was quenched with aq. $NaHCO_3$, extracted with EtOAc (2x), the organic phases collected, dried over $Na_2SO_4$, and then concentrated to dryness. The crude product was purified by automated silica gel chromatography to give a compound **3.1** as a yellow oil (1.2 g, 80%): LCMS [M+H] 293.0, ≥90% (214, 254 nm).

*Step 2. Tert-butyl ((2-((trimethylsilyl)ethynyl)thiazol-4-yl)methyl)carbamate (**3.2**).*

To a solution of *tert*-butyl ((2-bromothiazol-4-yl)methyl)carbamate (500 mg, 1.7 mmol) in 10 ml DMF:THF=1:1, were added $Pd(PPh_3)Cl_2$ (60 mg, 0.09 mmol), TEA (519 mg, 5.1 mmol), CuI (33 mg, 0.2 mmol), ethynyltrimethylsilane (252 mg, 2.6 mmol). The mixture was stirred at 60°C for 16h under $N_2$. The mixture was extracted with EtOAc (2x), the organic phases collected, dried over $Na_2SO_4$, and concentrated to dryness. The crude product was purified by automated silica gel chromatography to give compound **3.2** as a dark solid (276 mg, 52%): LCMS [M+H] 311.1, ≥90% (214, 254 nm).

*Step 3. (2-((Trimethylsilyl)ethynyl)thiazol-4-yl)methanamine (**3.3**).*

To a solution of *tert*-butyl ((2-((trimethylsilyl)ethynyl)thiazol-4-yl)methyl)carbamate (168 mg, 0.5 mmol) in DCM (2 mL), were added TBDMOTF( 0.5 mL) at rt for 1h. The mixture was basified with saturated aqueous $Na_2CO_3$ to pH 9 and diluted with MeOH and EtOAc. The organic phase was concentrated to obtain compound **3.3** (88 mg, 78%) as a yellow oil which was used directly in step 4 without further purification: LCMS [M+H] 211.1, ≥90% (214, 254 nm).

*Step 4. N-((2-((Trimethylsilyl)ethynyl)thiazol-4-yl)methyl)-1H-pyrazole-3-carboxamide (**3.4**).*

To a solution of (2-((trimethylsilyl)ethynyl)thiazol-4-yl)methanamine (88 mg, 0.4 mmol) in 5 mL DCM was added TEA (127 mg, 1.3 mmol) at rt. The mixture was cooled to 0$^o$C and 1*H*-pyrazole-3-carbonyl chloride (54 mg, 0.4 mmol) was added. The mixture was allowed to warm to rt and stir for 16h under $N_2$. To the reaction $H_2O$ (5 mL) was added and then diluted with DCM. The organic phase was dried with $Na_2SO_4$ and the crude product was purified by automated flash chromatography to give compound **3.4** as a white solid (20 mg, 16 %): LCMS [M+H] 305.1, ≥95% (214, 254 nm).

*Step 5. N-((2-Ethynylthiazol-4-yl)methyl)-1H-pyrazole-3-carboxamide (**4**).*

To a solution of *N*-((2-((trimethylsilyl)ethynyl)thiazol-4-yl)methyl)-1*H*-pyrazole-3-carboxamide (20 mg, 0.07 mmol) in 0.5 mL THF were added TBAF·THF (0.1 mL, 0.1 mmol) at rt for 1h. The mixture was purified by RP-HPLC to give final title compound **4** as a white solid (7 mg, 8%): LCMS [M+H] 233.1, HPLC ≥95% (254 nm); $^1$H NMR (400 MHz, DMSO) δ 13.27 (s, 1H), 8.67 (s, 1H), 7.82 (s, 1H), 7.48 (s, 1H), 6.66 (s, 1H), 4.89 (s, 1H), 4.52 (d, J=6.0 Hz, 2H).

***NMR***

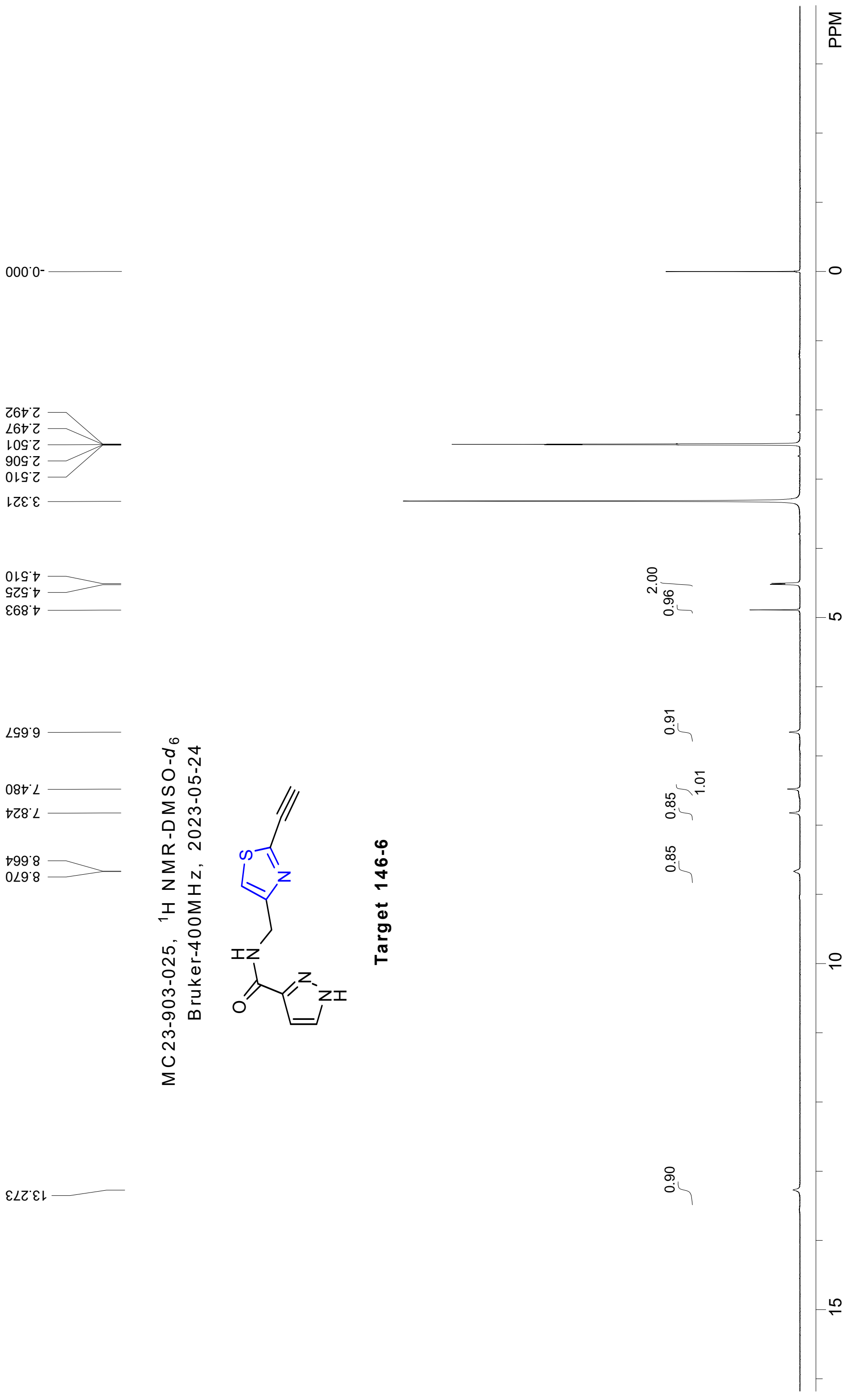

MC23-903-025, 1H NMR-DMSO-d6
Bruker-400MHz, 2023-05-24
Target 146-6
PPM
-0.000
2.492
2.497
2.501
2.506
2.510
3.321
4.510
4.525
4.893
6.657
7.480
7.824
8.664
8.670
13.273
2.00
0.96
0.91
1.01
0.85
0.85
0.90
0
5
10
15

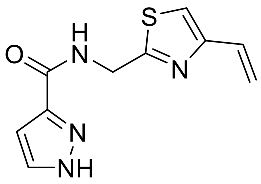

**_N_-((4-Vinylthiazol-2-yl)methyl)-1_H_-pyrazole-3-carboxamide (5).**

To a solution of intermediate **2.3** _N_-((4-bromothiazol-2-yl)methyl)-1_H_-pyrazole-3-carboxamide (100 mg, 0.4 mmol, see Scheme 2 above) in 3.5 mL of degassed toluene was added tributyl(vinyl)stannane (0.3 mL, 1.1 mmol) and $Pd(PPh_3)_4$. The mixture was stirred at $100^{o}C$ for 16h under $N_2$. The reaction mixture was cooled to rt and 2 mL $H_2O$ added. The mixture was extracted with EtOAc (2 x 5 mL), the organic phases washed with brine and dried over $Na_2SO_4$. After solvent removal under reduced pressure the crude material was dissolved in DMF and purified by RP-HPLC to give a white solid (9.6 mg, 12% yield): LCMS [M+H] 235.2; HPLC ≥95% (254 nm); $^{1}H$ NMR (400 MHz, DMSO-$d_6$) δ = 9.07 (s, 1H), 7.81 (s, 1H), 7.60 (d, _J_=99.0, 1H), 6.72 (dd, _J_=17.3, 10.9, 2H), 5.93 (dd, _J_=17.3, 2.0, 1H), 5.31 (dd, _J_=10.8, 2.0, 1H), 4.67 (d, _J_=6.2, 2H).

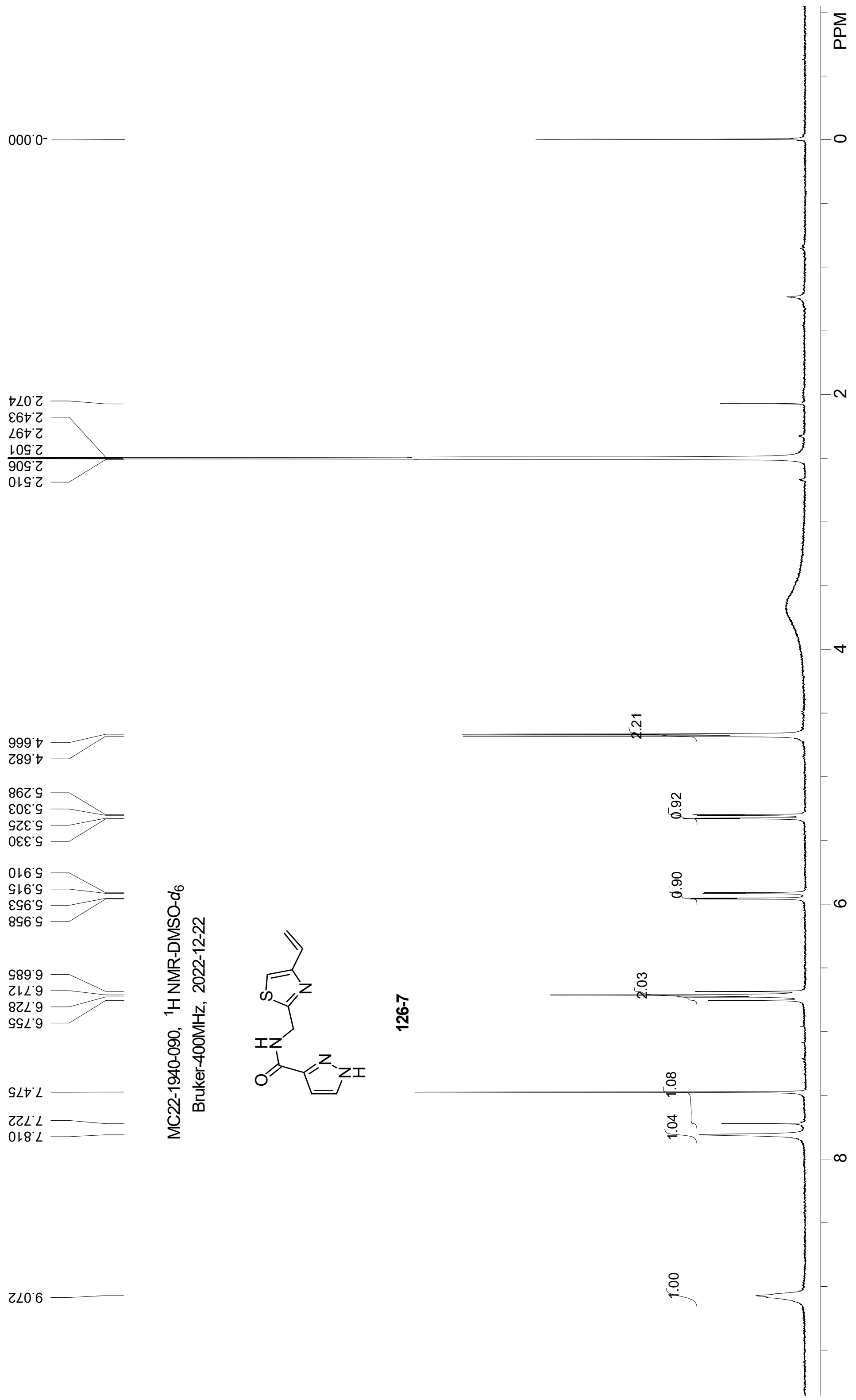
MC22-1940-090, 1H NMR-DMSO-d6
Bruker-400MHz, 2022-12-22
126-7
PPM
0
2
4
6
8
-0.000
2.074
2.493
2.497
2.501
2.506
2.510
4.666
4.682
5.298
5.303
5.325
5.330
5.910
5.915
5.953
5.958
6.685
6.712
6.728
6.755
7.475
7.722
7.810
9.072
2.21
0.92
0.90
2.03
1.08
1.04
1.00

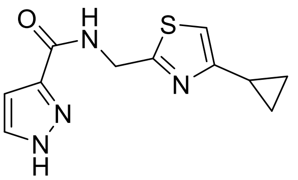

**N-((4-Cyclopropylthiazol-2-yl)methyl)-1*H*-pyrazole-3-carboxamide (6).**

***Batch 1 Synthesis***

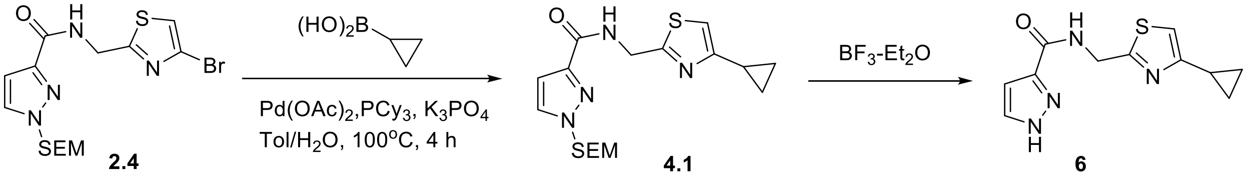


**Scheme 4.** Synthesis of compound **6** from intermediate **2.4**.

*Step 1. N-((4-Cyclopropylthiazol-2-yl)methyl)-1-((2-(trimethylsilyl) ethoxy)methyl)-1H-pyrazole-3-carboxamide (**4.1**)*

A 20 mL reaction vial was charged with *N*-((4-bromothiazol-2-yl)methyl)-1-((2-(trimethylsilyl)ethoxy)methyl)-1*H*-pyrazole-3-carboxamide (360 mg, 0.863 mmol), cyclopropylboronic acid (148 mg, 1.72 mmol), $K_2CO_3$ (380 mg, 1.72 mmol), $Pd(OAc)_2$ (28 mg, 0.17 mmol), and $PCy_3$ (48 mg, 0.17 mmol). A degassed solution of toluene/$H_2O$ (10 mL / 2 mL) was added and the mixture heated at 100°C for 4h under an atmosphere of $N_2$. The mixture was extracted with EtOAc (2 x 10 mL), the organic phases washed with brine and dried over $Na_2SO_4$. The filtrate was concentrated under reduced pressure and the crude material purified by automated silica gel chromatography to give *N*-((4-cyclopropylthiazol-2-yl)methyl)-1-((2-(trimethylsilyl)ethoxy)methyl)-1*H*-pyrazole-3-carboxamide (290 mg, 89% yield): LCMS [M+H]: 379.2, ≥95% (214, 254 nm).

*Step 2. N-((4-Cyclopropylthiazol-2-yl)methyl)-1H-pyrazole-3-carboxamide (**6**).*

To a solution of *N*-((4-cyclopropylthiazol-2-yl)methyl) -1-((2-(trimethylsilyl) ethoxy)methyl)-1*H* -pyrazole-3-carboxamide (290 mg, 0.8 mmol) in DCM (10 mL) was added $BF_3$-$Et_2O$ (0.6 mL) dropwise. The mixture was stirred at rt for 1h and then $NH_3$-$H_2O$ (1.8 mL) added.

The mixture was stirred for an additional 18h, extracted with EtOAc (2x), and the organic layers combined and dried over $Na_2SO_4$. The filtrate was concentrated and purified by RP-HPLC to give title compound **6** (6.5 mg): LCMS [M+H] 249.1, HPLC ≥95% (254 nm); $^1$H NMR (400 MHz, MeOD) δ 7.73 (s, 1H), 6.99 (s, 1H), 6.81 (s, 1H), 4.78 (s, 2H), 2.10 – 1.98 (m, 1H), 0.98 – 0.89 (m, 2H), 0.86 – 0.77 (m, 2H).

***Batch 2 Synthesis 6.***

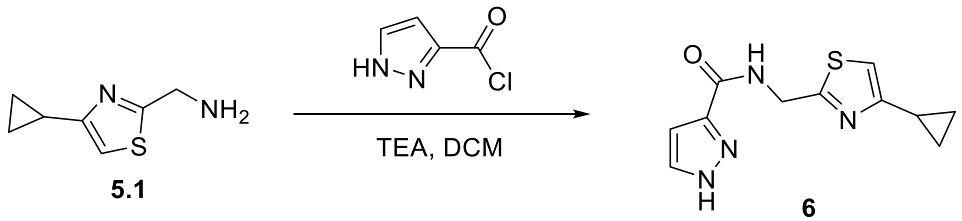


**Scheme 5.** Batch 2 direct synthesis of **6** using compound **5.1**.

**Preparation of (4-cyclopropylthiazol-2-yl)methanamine (5.1) from ethyl 4-cyclopropylthiazole-2-carboxylate.**

Amine **5.1** was synthesized three steps starting from commercial ethyl 4-cyclopropylthiazole-2-carboxylate using a protocol adopted from WO2023001061:

*Step 1*. A mixture of ethyl 4-cyclopropylthiazole-2-carboxylic acid (2.5 g, 12.7 mmol, CAS 1083274-67-2) in $NH_3$/MeOH (20 mL) was stirred at rt for 18h. The mixture was concentrated to afford 4-cyclopropylthiazole-2-carboxamide (2.1 g, 98% yield): LCMS [M+H] 169.1.

*Step 2*. To a mixture of 4-cyclopropylthiazole-2-carboxamide (3 .0 g, 17.9 mmol) in DCM (20 mL) was added TEA (6.2 mL, 44.6 mmol) and TFAA (6.2 mL, 44.6 mmol) at 0$^o$C. The mixture warmed to rt and stirred for 18h. The mixture was concentrated and the crude product purified by automated silica gel chromatography (EtOAc/hexanes) to give 4-cyclopropylthiazole-2-carbonitrile (2.6 g,  96% yield): LCMS [M+H] 151.1.

*Step 3*. To 4-cyclopropylthiazole-2-carbonitrile (2.7 g, 18.2 mmol) in THF (30 mL) was added Raney Ni (2.0 g) at 0$^{o}$C. The mixture warmed to rt and stirred for 18h. The mixture was then filtered over Celite, concentrated, and purified using a short pad of silica gel (MeOH/DCM) to give desired **5.1** (4-cyclopropylthiazol-2-yl)methanamine (2.8 g, 98% yield): LCMS [M+H]:155.0, ≥93% (214, 254 nm).

***N*-((4-Cyclopropylthiazol-2-yl)methyl)-1*H*-pyrazole-3-carboxamide (6).**

A 20 mL reaction vial was charged with **5.1** (4-cyclopropylthiazol-2-yl)methanamine (300 mg, 1.95 mmol) in DCM (5 mL) and TEA (0.54 mL, 1.56 mmol). The mixture was cooled to 0$^{o}$C and 1*H*-pyrazole-3-carbonyl chloride (203 mg, 1.56 mmol) was added. After 2h the mixture warmed to rt and concentrated to afford crude product. The mixture was purified by RP-HPLC to give *N*-((4-cyclopropylthiazol-2-yl)methyl)-1*H*-pyrazole-3-carboxamide (303 mg, 63%): LCMS [M+H] 249.1; HPLC ≥95% (254 nm); $^{1}$H NMR (400 MHz, DMSO) δ 13.30 (s, 1H), 8.95 (s, 1H), 7.84 (s, 1H), 7.11 (s, 1H), 6.67 (s, 1H), 4.60 (d, *J* = 5.9 Hz, 2H), 2.03 (td, *J* = 8.2, 4.1 Hz, 1H), 0.91 – 0.82 (m, 2H), 0.81 – 0.75 (m, 2H).

368 ***Batch 1 NMR***

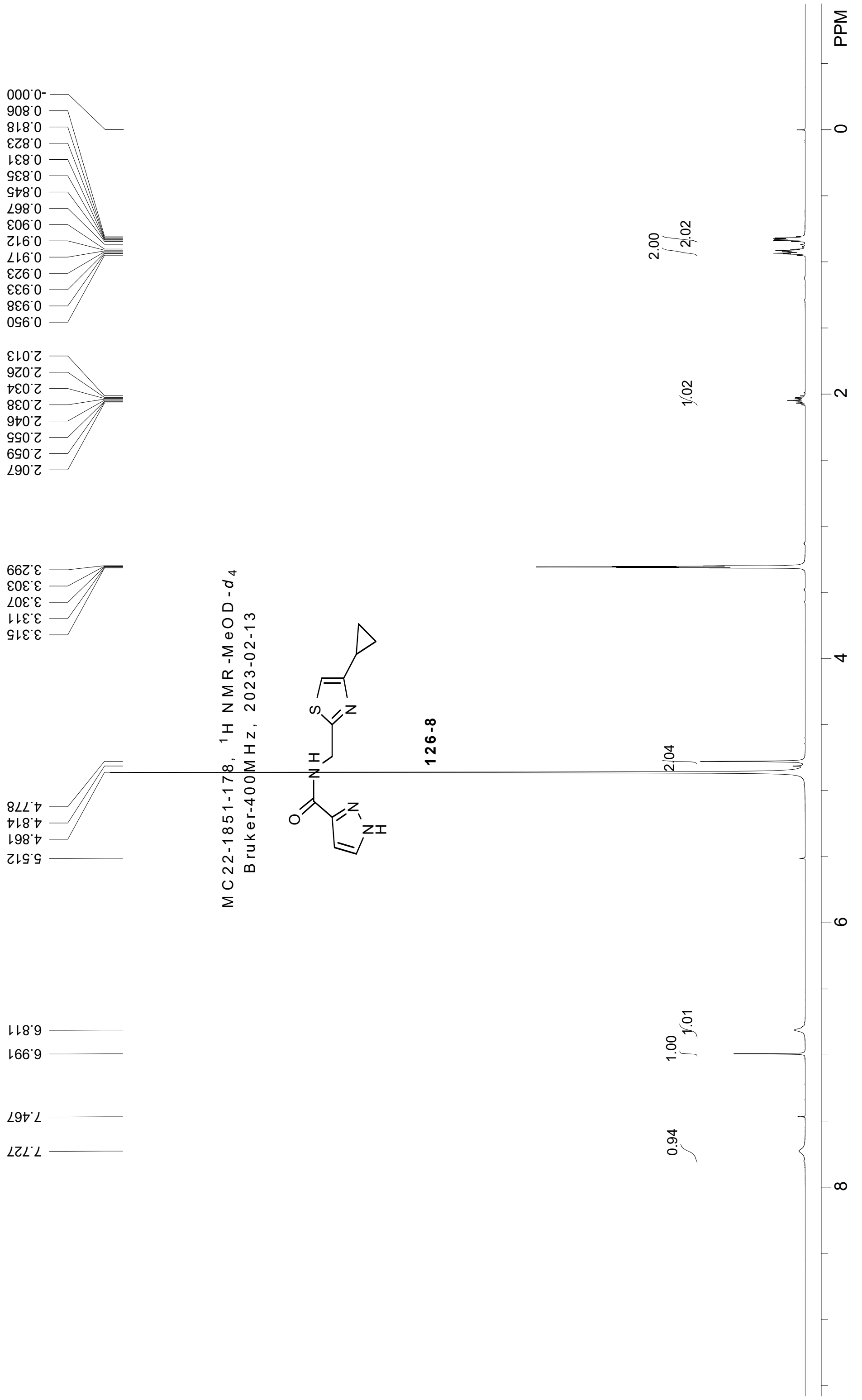


369

### *Batch 2 NMR*

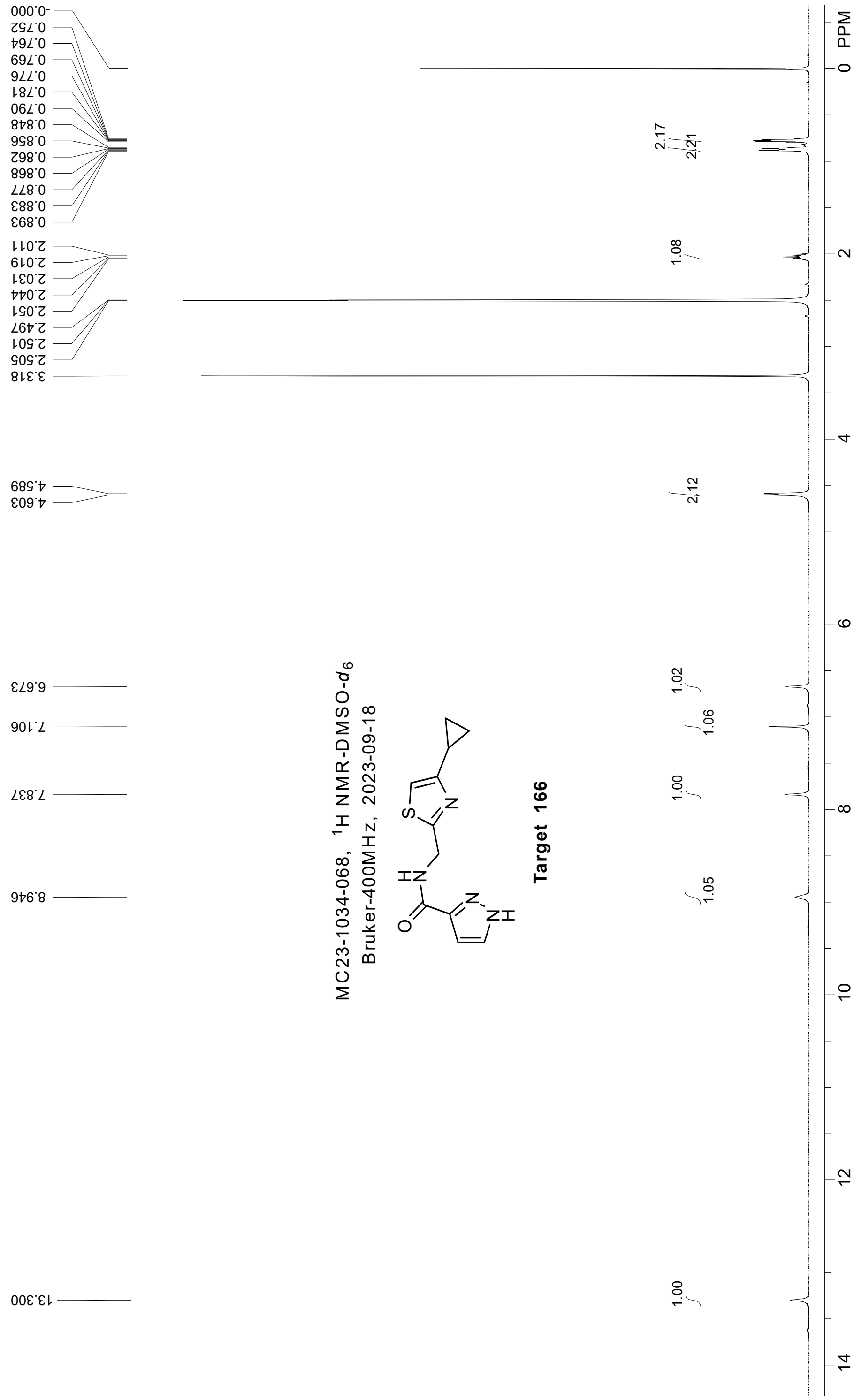

MC23-1034-068, 1H NMR-DMSO-d6
Bruker-400MHz, 2023-09-18
Target 166
PPM
-0.000
0.752
0.764
0.769
0.776
0.781
0.790
0.848
0.856
0.862
0.868
0.877
0.883
0.893
2.011
2.019
2.031
2.044
2.051
2.497
2.501
2.505
3.318
4.589
4.603
6.673
7.106
7.837
8.946
13.300
2.17
2.21
1.08
2.12
1.02
1.06
1.00
1.05
1.00

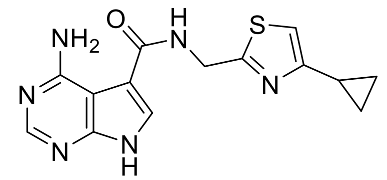

**4-Amino-*N*-((4-cyclopropylthiazol-2-yl)methyl)-7*H*-pyrrolo[2,3-*d*]pyrimidine-5-carboxamide (7).**

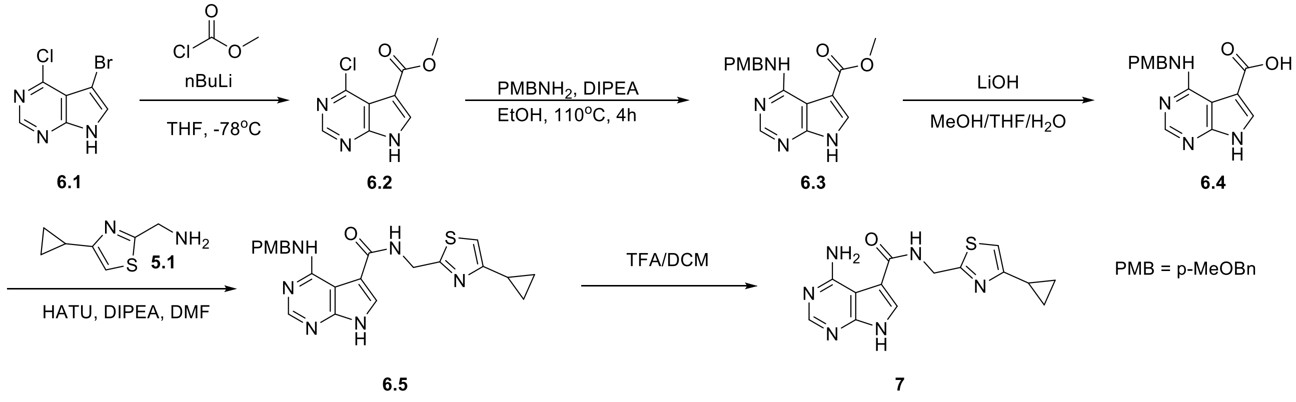

**Scheme 6.** Synthesis of merger target **7**.

*Step 1. Methyl 4-chloro-7H-pyrrolo[2,3-d]pyrimidine-5-carboxylate (**6.2**).*

To a -78°C solution of 5-bromo-4-chloro-7(*H)*-pyrrolo[2,3-*d*]pyrimidine (500 mg, 2.15 mmol, CAS 22276-95-5) and methyl carbonochloridate (194 mg, 2.05 mmol) in THF (10 mL) under $N_2$ was added *n*-BuLi (2.7 mL, 4.30 mmol) dropwise. The mixture was stirred at -78°C for 30 min, then rt for 3h. The mixture was slowly quenched with aqueous $NH_4Cl$ at rt and extracted with EtOAc (2x). The combined organic layers were washed with brine, dried over $Na_2SO_4$, and concentrated under reduced pressure. The crude product was purified by automated silica gel chromatography (EtOAc/hexanes) to give methyl 4-chloro-7*H*-pyrrolo[2,3-*d*]pyrimidine-5-carboxylate (269 mg, 62%): LCMS [M+H] 212.1, ≥95% (214, 254 nm).

*Step 2. Methyl 4-((4-methoxybenzyl)amino)-7H-pyrrolo[2,3-d]pyrimidine-5-carboxylate (**6.3**).*

To a solution of methyl 4-chloro-7*H*-pyrrolo[2,3-*d*]pyrimidine-5-carboxylate (220 mg, 1.04 mmol) in EtOH (10 mL) at 110°C was added DIPEA (268 mg, 2.08 mmol) and 4-

methoxybenzyl amine (143 mg, 1.04 mmol) under $N_2$. The mixture was stirred at 110°C for 3h. The mixture was concentrated to afford a crude product to give methyl 4-((4-methoxybenzyl)amino)-7*H*-pyrrolo[2,3-*d*]pyrimidine-5-carboxylate (230 mg, 71%). LCMS [M+H]: 313.1, ≥95% (214, 254 nm).

*Step 3. 4-((4-Methoxybenzyl)amino)-7H-pyrrolo[2,3-d]pyrimidine-5-carboxylic acid (**6.4**).*

To a solution of methyl **6.3** 4-((4-methoxybenzyl)amino)-7*H*-pyrrolo[2,3-*d*]pyrimidine-5-carboxylate (230 mg, 0.74 mmol) in MeOH (3 mL), THF (3 mL) and $H_2O$ (3 mL), at rt was added NaOH (590 mg, 14.7 mmol). The mixture was warmed to 50°C and stirred for 18h. The mixture was acidified with 1N HCl to pH 3, extracted with EtOAc, dried over $Na_2SO_4$, and concentrated to give 4-((4-methoxybenzyl)amino)-7*H*-pyrrolo[2,3-*d*]pyrimidine-5-carboxylic acid (135 mg, 61%): LCMS [M+H] 299.0, ≥90% (214, 254 nm).

*Step 4. N-((4-Cyclopropylthiazol-2-yl)methyl)-4-((4-methoxybenzyl)amino)-7H-pyrrolo[2,3-d]pyrimidine-5-carboxamide (**6.5**).*

To a 0°C solution of 4-((4-methoxybenzyl)amino)-7*H*-pyrrolo[2,3-*d*]pyrimidine-5-carboxylic acid (135 mg, 0.45 mmol, intermediate **5.1** synthesis described above for **6** Batch 2, above) in DMF (5 mL) was added DIPEA (116 mg, 0.90 mmol) and HATU (256 mg, 0.68 mmol) under $N_2$. The mixture was stirred at 0°C for 15 min and (4-cyclopropylthiazol-2-yl)methanamine (77 mg, 0.50 mmol, 1.1 eq) added. The mixture warmed to rt and stirred for 18h. The mixture was diluted with EtOAc, the organic layer separated, washed with brine, then dried over $Na_2SO_4$ and concentrated. The crude product was purified by automated silica gel chromatography (EtOAc/hexanes) to give **6.5** *N*-((4-cyclopropylthiazol-2-yl)methyl)-4-((4-methoxybenzyl)amino)-7*H*-pyrrolo[2,3-d]pyrimidine-5-carboxamide (120 mg, 61%): LCMS [M+H] 435.1, ≥90% (214, 254 nm).

*Step 5. 4-Amino-N-((4-cyclopropylthiazol-2-yl)methyl)-7H-pyrrolo[2,3-d]pyrimidine-5-carboxamide (**7**).*

To mixture of *N*-((4-cyclopropylthiazol-2-yl)methyl)-4-((4-methoxybenzyl)amino)-7*H*-pyrrolo[2,3-*d*]pyrimidine-5-carboxamide (80 mg, 0.16 mmol) in TFA (5.0 mL) was stirred at 50°C for 6h. The mixture was concentrated to afford crude material which purified by

RP-HPLC to give **7** 4-amino-*N*-((4-cyclopropylthiazol-2-yl)methyl)-7*H*-pyrrolo[2,3-*d*]pyrimidine-5-carboxamide (30 mg, 60%): LCMS [M+H] 315.1; HPLC ≥95% (254 nm); $^{1}$H NMR (400 MHz, DMSO-*d*6) δ 12.47 (s, 1H), 9.32 (s, 1H), 8.15 (d, *J* = 14.2 Hz, 1H), 8.06 (s, 1H), 4.67 (d, *J* = 6.0 Hz, 2H), 2.10 – 1.99 (m, 1H), 0.97 – 0.83 (m, 2H), 0.82 – 0.68 (m, 2H).

***NMR***

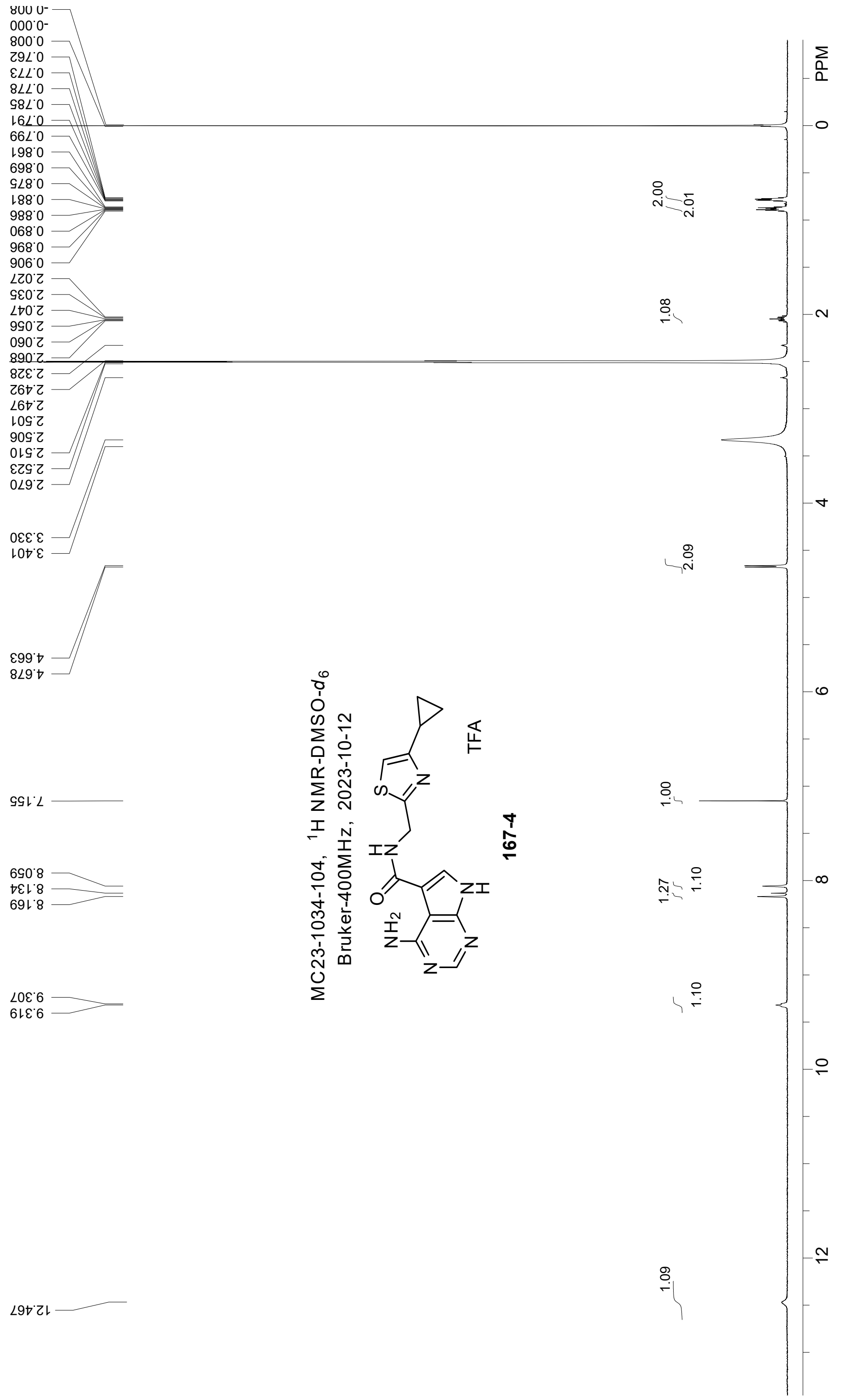

MC23-1034-104, 1H NMR-DMSO-d6
Bruker-400MHz, 2023-10-12
TFA
167-4
PPM

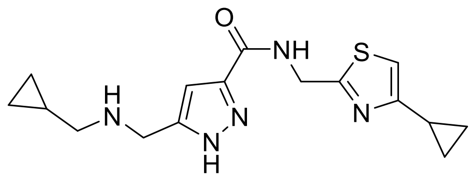

**5-(((Cyclopropylmethyl)amino)methyl)-N-((4-cyclopropylthiazol-2-yl)methyl)-1H-pyrazole-3-carboxamide (12).**

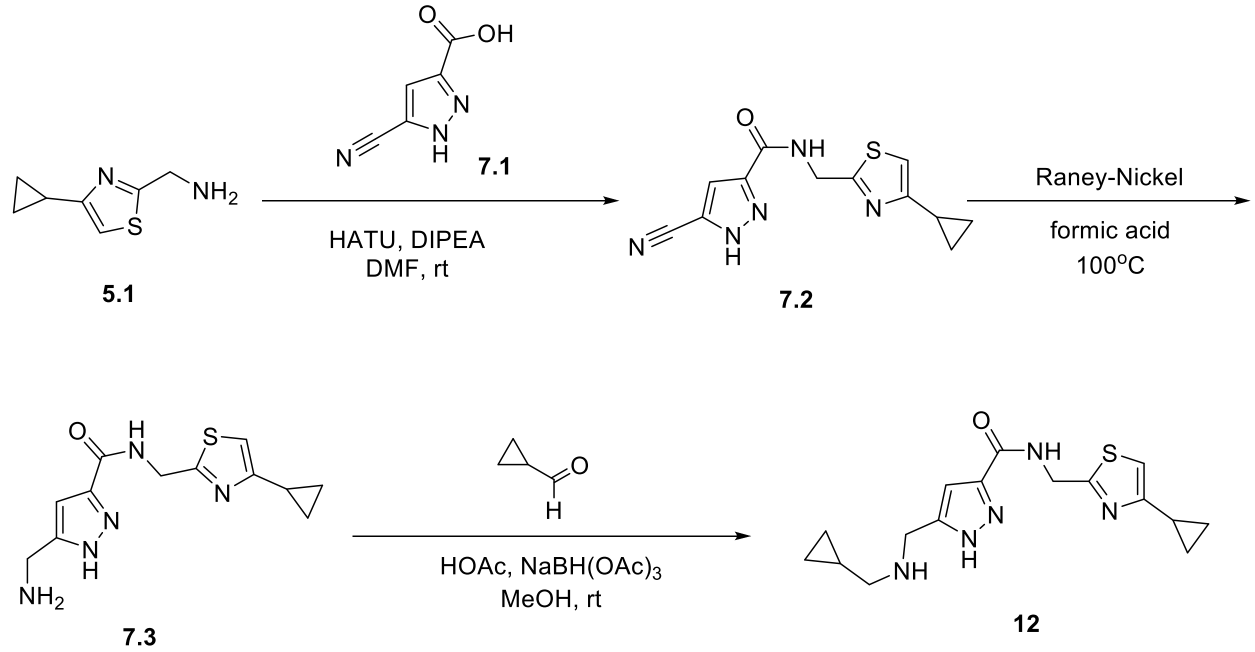

**Scheme 7.** Synthesis route for merger target **12** (batch 1 and 2).

Step 1. 5-Cyano-*N*-((4-cyclopropylthiazol-2-yl)methyl)-1*H*-pyrazole-3-carboxamide (**7.2**). A mixture of compound **5.1** (100 mg, 0.730 mmol), commercially available 5-cyano-1*H*-pyrazole-3-carboxylic acid **7.1** (124 mg, 0.80 mmol, CAS 1187361-13-2), HATU (416 mg, 1.10 mmol) and DIPEA (189 mg, 1.46 mmol) in DMF (5 mL) was stirred at rt for 6h under an inert atmosphere. The mixture was extracted with EtOAc (2x) and the combined extracts washed with brine. The combined organic layers were dried over $Na_2SO_4$ and concentrated. The crude was purified by automated silica gel chromatography (EtOAc/hexanes) to afford **7.2** as a brown solid (105 mg, 53%): LCMS [M+H] 274.0, ≥95% (214, 254 nm).

Step 2. 5-(Aminomethyl)-*N*-((4-cyclopropylthiazol-2-yl)methyl)-1*H*-pyrazole-3-carboxamide (**7.3**).

A mixture of compound **7.2** (95 mg, 0.35 mmol) and Raney-Nickel (36 mg) in formic acid (5 mL) was stirred at 100°C for 0.5h. LCMS showed the reaction was completed. The mixture was purified by RP-HPLC. The title compound **7.3** was formed as a white solid (77 mg, 83%): LCMS [M+H] 278.1, ≥95% (214, 254 nm); $^{1}H$ NMR (400 MHz, DMSO) δ 5.90 (s, 2H), 5.05 (s, 1H), 4.60 (s, 1H), 2.67 (s, 3H), 1.95 (s, 2H), -0.25 (s, 1H), -1.27 (d, J = 7.2 Hz, 2H), -1.50 (s, 2H).

*Step 3. 5-(((Cyclopropylmethyl)amino)methyl)-N-((4-cyclopropylthiazol-2-yl)methyl)-1H-pyrazole-3-carboxamide* (**12**).

To a mixture of **7.3** 5-(aminomethyl)-*N*-((4-cyclopropylthiazol-2-yl)methyl)-1*H*-pyrazole-3-carboxamide (60 mg, 0.22 mmol) in MeOH (5 mL) was added cyclopropanecarbaldehyde (15.4 mg, 0.22 mmol), HOAc (one drop) and $NaHB(OAc)_3$ (187 mg, 0.88 mmol). The mixture was stirred at rt for 5h, diluted with EtOAc and water. The organic phase was washed with brine, dried over $Na_2SO_4$, and concentrated to dryness. The crude material was purified by RP-HPLC to afford title compound **12** as a white solid (6 mg, 10%). Batch 2 was synthesized by the same protocol on larger scale (200 mg of **7.3**, 51 mg product obtained): LCMS [M+H] 332.2, ≥95% (214, 254 nm); HPLC ≥95% (254 nm); $^{1}H$ NMR (400 MHz, DMSO) δ 13.94 (s, 1H), 9.43 (s, 1H), 9.03 (s, 2H), 7.15 (s, 1H), 6.94 (s, 1H), 4.64 (d, J = 5.2 Hz, 2H), 4.22 (s, 2H), 2.87 (d, J = 5.2 Hz, 2H), 2.13 – 1.95 (m, 1H), 1.05 (s, 1H), 0.88 (m, 2H), 0.80 – 0.73 (m, 2H), 0.58 (m, 2H), 0.35 (d, J = 6.0 Hz, 2H).

### ***Batch 1 NMR***

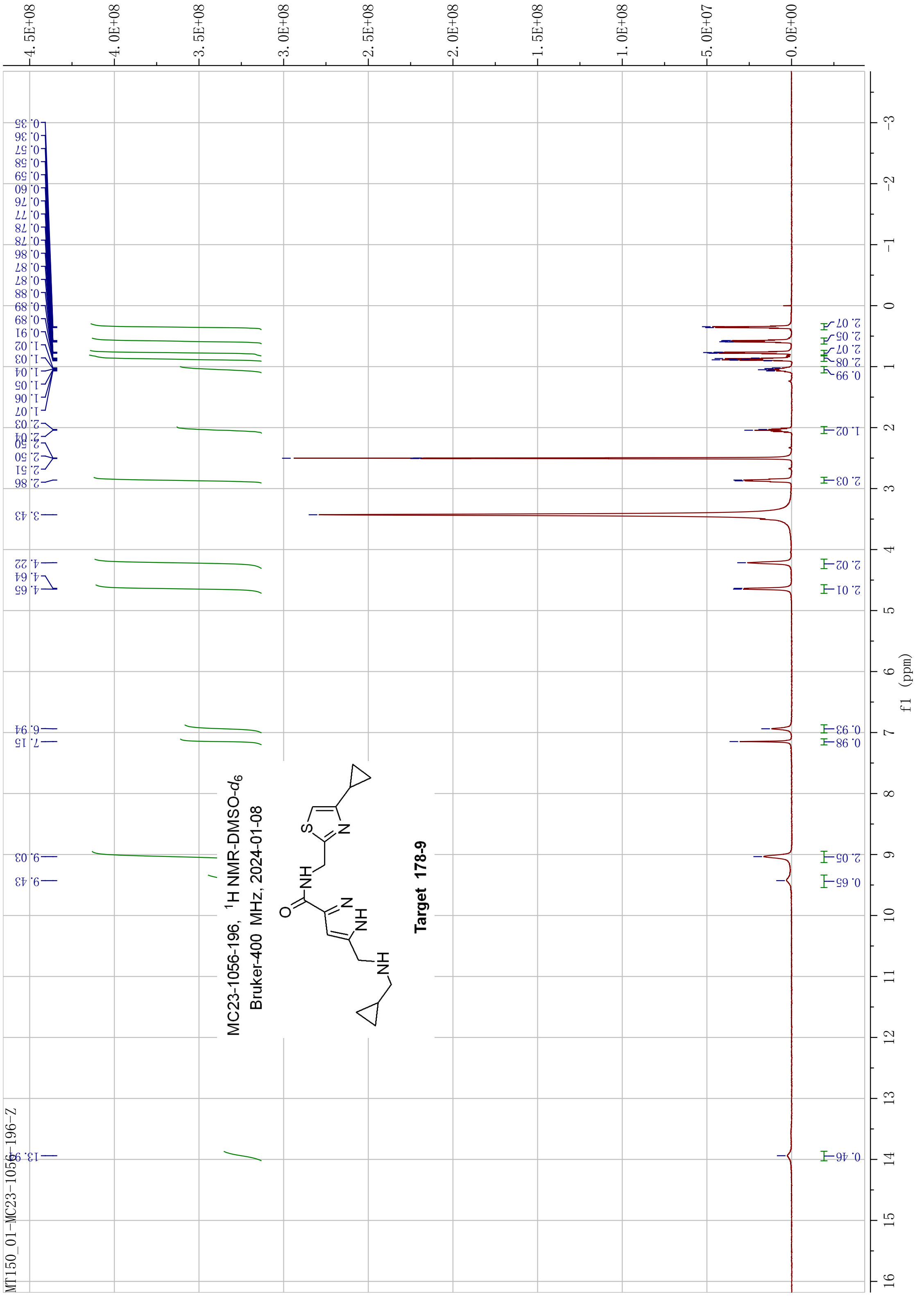
MC23-1056-196, 1H NMR-DMSO-d6
Bruker-400 MHz, 2024-01-08
Target 178-9
MT150_01-MC23-1056-196-Z
f1 (ppm)

### *Batch 2 NMR*

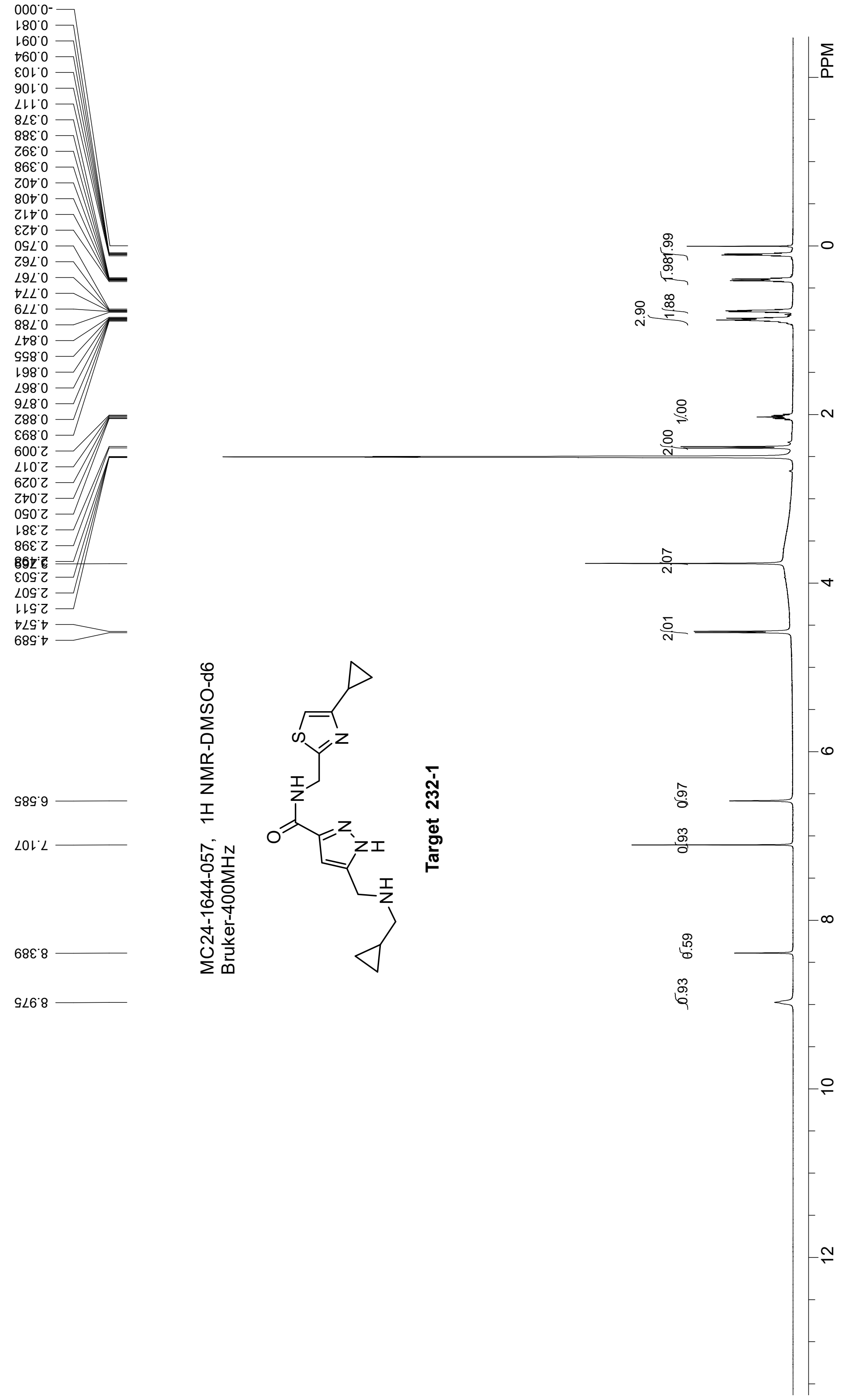

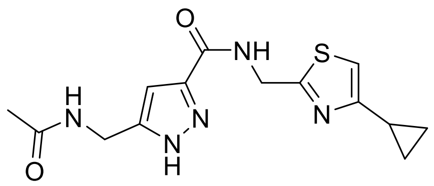

**5-(Acetamidomethyl)-*N*-((4-cyclopropylthiazol-2-yl)methyl)-1*H*-pyrazole-3-carboxamide (13).**

A mixture of compound **7.3** (100 mg, 0.36 mmol, synthesized as described above) and $Ac_2O$ (37 mg, 0.36 mmol) in HOAc (5 mL) was stirred at rt for 1h. The mixture was concentrated and purified by RP-HPLC to obtain title compound **13** as a white solid (4 mg, <5%). Batch 2 was produced using the same synthesis and purification protocol on larger scale (295 mg, 31 mg product): LCMS [M+H], 320.1, ≥95% purity (214, 254 nm); $^1$H NMR (400 MHz, DMSO) δ 9.06 (t, J = 5.9 Hz, 1H), 8.37 (s, 1H), 7.13 (s, 1H), 6.59 (s, 1H), 4.59 (d, J = 6.2 Hz, 2H), 4.27 (d, J = 4.4 Hz, 2H), 2.10 – 1.98 (m, 1H), 1.86 (s, 3H), 0.96 – 0.83 (m, 2H), 0.80 – 0.73 (m, 2H).

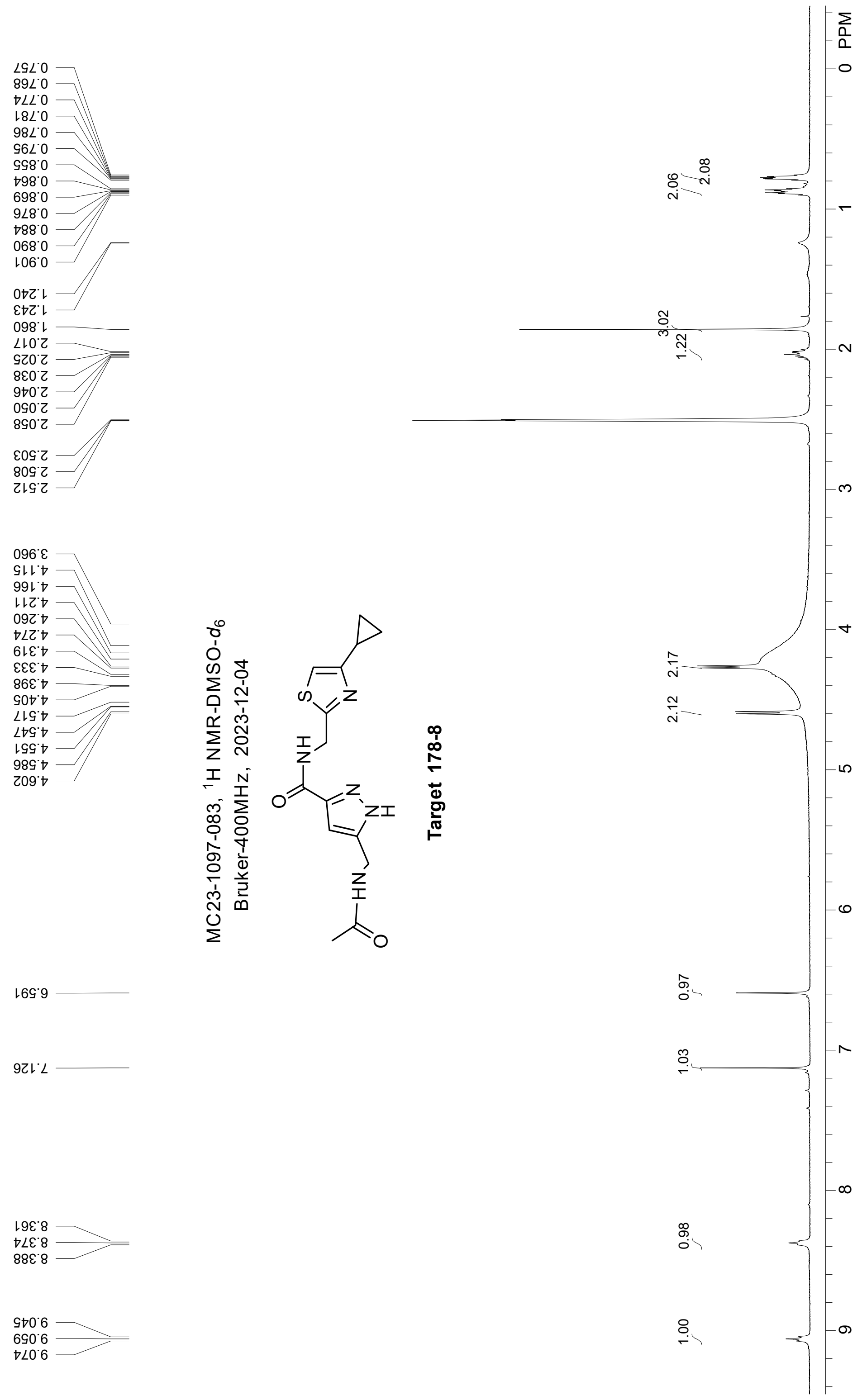
MC23-1097-083, 1H NMR-DMSO-d6
Bruker-400MHz, 2023-12-04
Target 178-8
0.757
0.768
0.774
0.781
0.786
0.795
0.855
0.864
0.869
0.876
0.884
0.890
0.901
1.240
1.243
1.860
2.017
2.025
2.038
2.046
2.050
2.058
2.503
2.508
2.512
3.960
4.115
4.166
4.211
4.260
4.274
4.319
4.333
4.398
4.405
4.517
4.547
4.551
4.586
4.602
6.591
7.126
8.361
8.374
8.388
9.045
9.059
9.074
2.06
2.08
3.02
1.22
2.17
2.12
0.97
1.03
0.98
1.00
PPM
0
1
2
3
4
5
6
7
8
9

### *Batch 2 NMR*

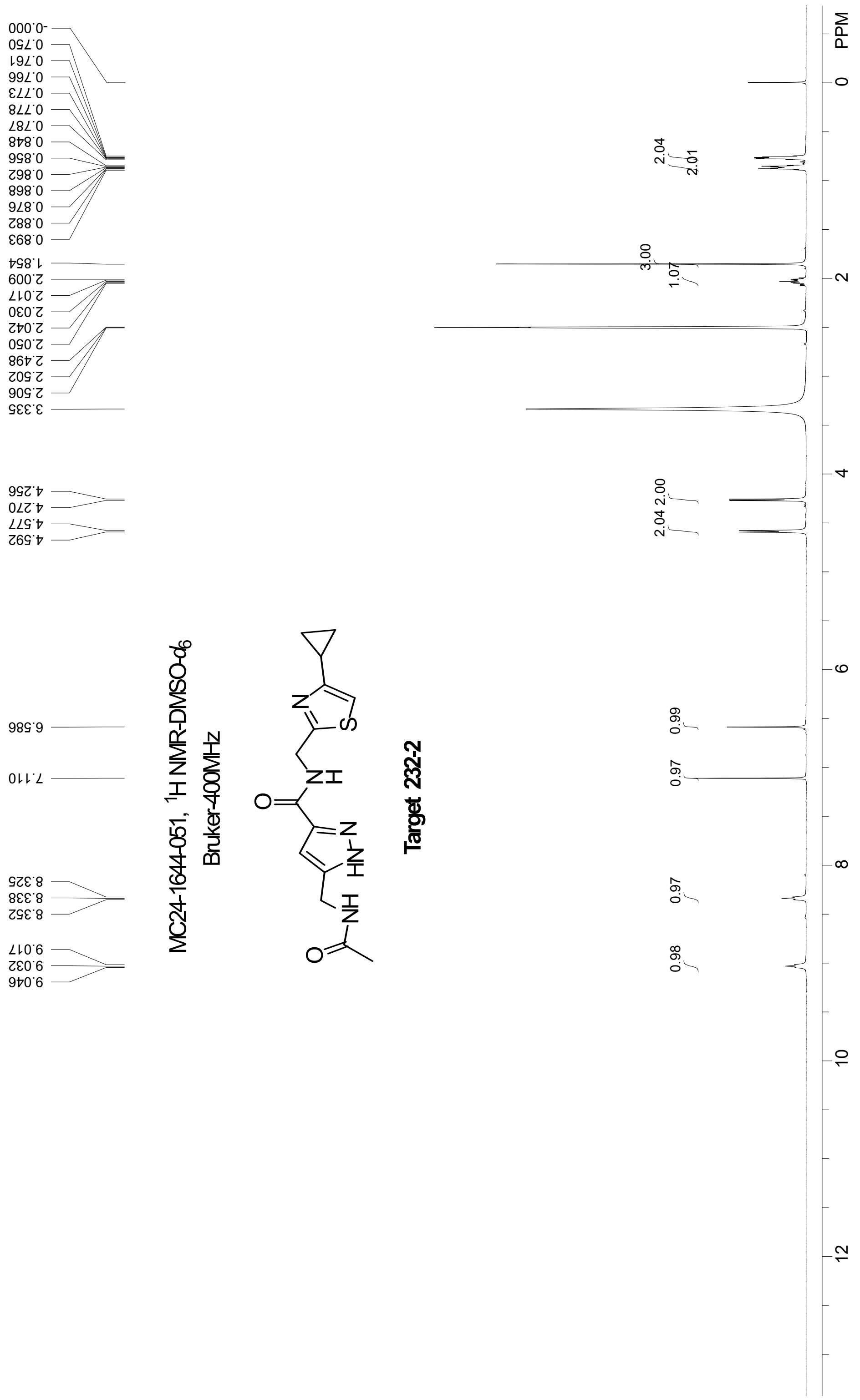
MC24-1644-051, ¹H NMR-DMSO-d6
Bruker-400MHz
Target 232-2
-0.000
0.750
0.761
0.766
0.773
0.778
0.787
0.848
0.856
0.862
0.868
0.876
0.882
0.893
1.854
2.009
2.017
2.030
2.042
2.050
2.498
2.502
2.506
3.335
4.256
4.270
4.577
4.592
6.586
7.110
8.325
8.338
8.352
9.017
9.032
9.046
2.04
2.01
3.00
1.07
2.00
2.04
0.99
0.97
0.97
0.98
PPM
0
2
4
6
8
10
12

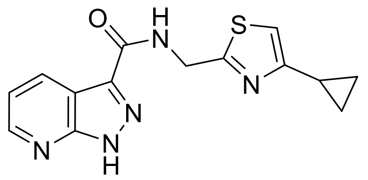


**N-((4-Cyclopropylthiazol-2-yl)methyl)-1*H*-pyrazolo[3,4-*b*]pyridine-3-carboxamide (14).**

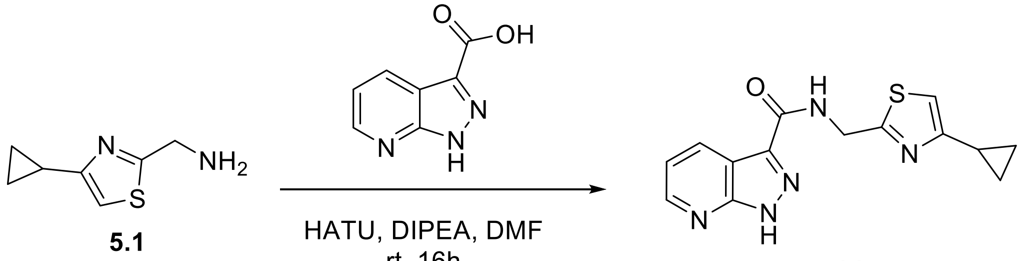


**Scheme 8.** Synthesis of merger target **14**.

To a solution of 1*H*-pyrazolo[3,4-b]pyridine-3-carboxylic acid (191 mg, 1.18 mmol, CAS 116855-08-4) in DMF (7 mL) was added DIPEA (304 mg, 2.36 mmol, 2.0 eq), HATU (673 mg, 1.77 mmol) at 0$^o$C under $N_2$. The mixture was stirred at 0$^o$C for 15 min, **5.1** (4-cyclopropylthiazol-2-yl)methanamine (200 mg, 1.30 mmol) was added, and then the reaction stirred at rt for 18h. The mixture was diluted with water, extracted with EtOAc (2x), the combined organic layers washed with brine, dried over $Na_2SO_4$, and then concentrated under reduced pressure. The crude was purified by RP-HPLC to give *N*-((4-cyclopropylthiazol-2-yl)methyl)-1*H*-pyrazolo[3,4-*b*]pyridine-3-carboxamide (83 mg, 24%): LCMS [M+H] 299.1; HPLC ≥95% (254 nm); $^1$H NMR (400 MHz, DMSO) δ 14.26 (s, 1H), 9.36 (t, *J* = 6.2 Hz, 1H), 8.61 (dd, *J* = 4.5, 1.6 Hz, 1H), 8.52 (dd, *J* = 8.0, 1.6 Hz, 1H), 7.35 (dd, *J* = 8.1, 4.4 Hz, 1H), 7.12 (s, 1H), 4.68 (d, *J* = 6.2 Hz, 2H), 2.15 – 1.91 (m, 1H), 0.92 – 0.85 (m, 2H), 0.83 – 0.74 (m, 2H).

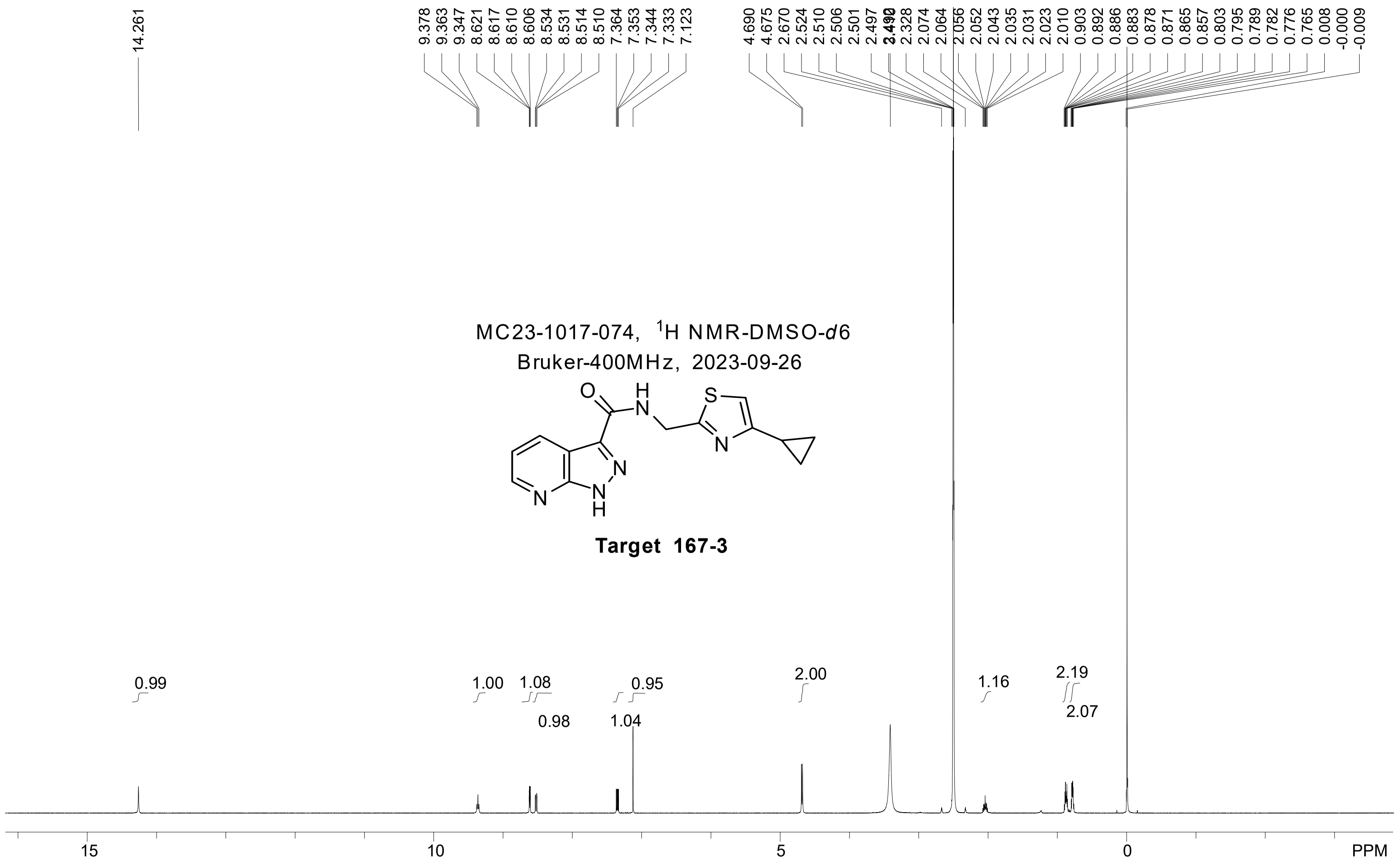
MC23-1017-074, 1H NMR-DMSO-d6
Bruker-400MHz, 2023-09-26
Target 167-3
14.261
9.378 9.363 9.347 8.621 8.617 8.610 8.606 8.534 8.531 8.514 8.510 7.364 7.353 7.344 7.333 7.123
4.690 4.675 2.670 2.524 2.510 2.506 2.501 2.497 2.490 2.328 2.074 2.064 2.056 2.052 2.043 2.035 2.031 2.023 2.010 0.903 0.892 0.886 0.883 0.878 0.871 0.865 0.857 0.803 0.795 0.789 0.782 0.776 0.765 0.008 -0.000 -0.009
0.99
1.00
1.08
0.98
0.95
1.04
2.00
1.16
2.19
2.07
15
10
5
0
PPM

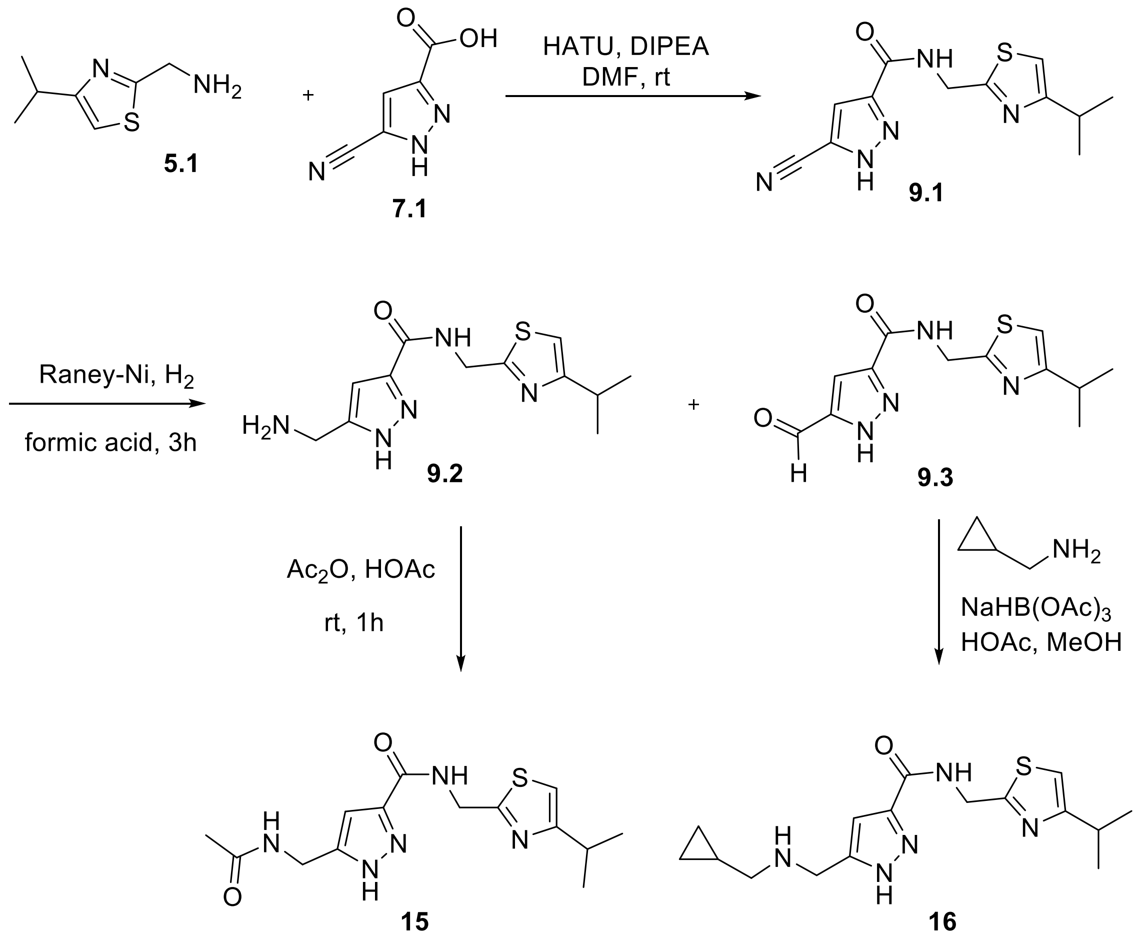


**Scheme 9.** Synthesis of merger targets **15** and **16**.

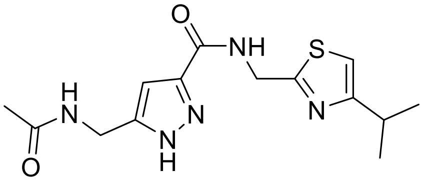

**5-(Acetamidomethyl)-N-((4-isopropylthiazol-2-yl)methyl)-1H-pyrazole-3-carboxamide (15).**

*Step 1. 5-Cyano-N-((4-isopropylthiazol-2-yl)methyl)-1H-pyrazole-3-carboxamide (**9.1**).* To a DMF (20 mL) solution of 5-cyano-1H-pyrazole-3-carboxylic acid (615 mg, 4.48 mmol) was added EDCI (1.03 g, 5.38 mmol), HOBT (727 mg, 5.38 mmol) and DIPEA (1.15 g, 8.9 mmol). The mixture was stirred at 0°C for 0.5h under an inert atmosphere and then (4-isopropylthiazol-2-yl)methanamine (700 mg,4.48 mmol) was added (amine **5.1**, was prepared according to metal-free conditions detailed above and in WO202300106). The mixture was stirred at rt for 16h. The mixture was quenched with $H_2O$ and extracted with EtOAc (2x). The combined organic layers were washed with brine, dried over $Na_2SO_4$, and concentrated to dryness. The crude product was purified by automated flash chromatography to give **9.1** as a white solid (370 mg, 30%): LCMS [M+H] 276.2, ≥90% (214, 254 nm).

*Step 2. 5-(Aminomethyl)-N-((4-isopropylthiazol-2-yl)methyl)-1H -pyrazole-3-carboxamide (**9.2**) and 5-formyl-N-((4-isopropylthiazol-2-yl)methyl)-1H-pyrazole -3-carboxamide (**9.3**).*

To a solution of **9.1** (220 mg, 0.8 mmol) in formic acid (5 mL) was added Raney-Nickel (50 mg). The mixture was stirred at rt for 3h under an atmospheric hydrogen balloon and by LCMS and TLC the reaction appeared to be complete and contain a ~1:1 mixture of **9.2** and **9.3**. The mixture was degassed and filtered over Celite. The filtrate was concentrated and purified by automated flash chromatography to give **9.2** as a yellow oil (80 mg, 35%) and **9.3** as a clear oil (74 mg, 31%). **9.2**: LCMS [M+H] 280.1, ≥93% (214, 254 nm); **9.3**: LCMS [M+H] 279.1, ≥90% (214, 254 nm).

*Step 3. 5-(Acetamidomethyl)-N-((4-isopropylthiazol-2-yl)methyl)-1H-pyrazole-3-carboxamide (**15**).*

To a solution of **9.2** 5-(aminomethyl)-*N*-((4-isopropylthiazol-2-yl)methyl)-1*H* -pyrazole-3-carboxamide (40 mg, 0.143 mmol) in AcOH (3 mL) was added $Ac_2O$ (12 mg, 0.114 mmol). The mixture was stirred at rt for 3h under inert atmosphere. The crude product was purified by PR-HPCL to give title compound **15** as a white solid (25 mg, 54%): LCMS [M+H] 322.2, ≥95% (214, 254 nm); HPLC ≥95% (254 nm); $^1$H NMR (400 MHz, MeOD) δ 7.03 (s, 1H), 6.66 (s, 1H), 4.79 (s, 2H), 4.42 (s, 2H), 3.15 – 2.93 (m, 1H), 1.98 (s, 3H), 1.29 (d, J = 7.0 Hz, 6H).

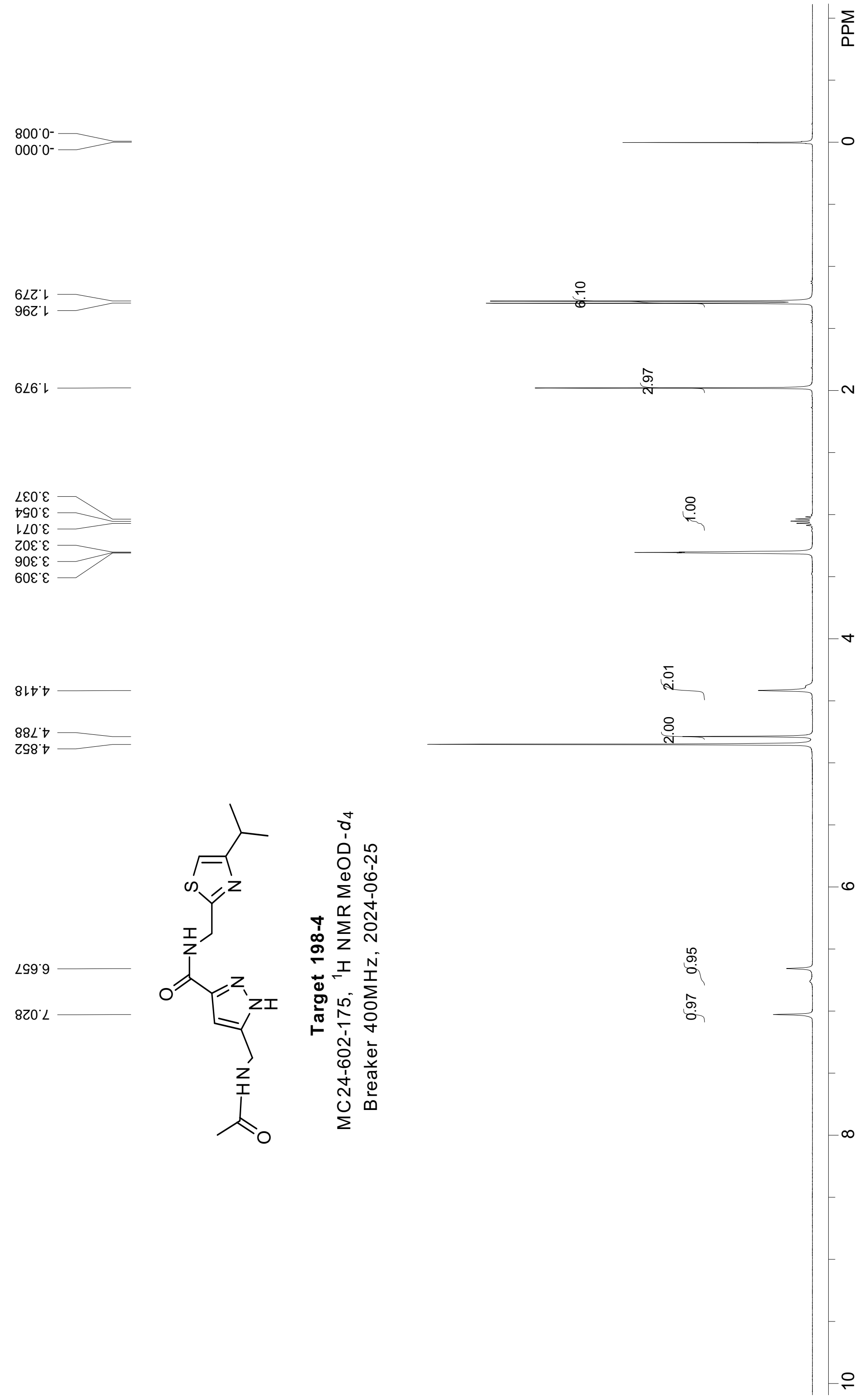


**Target 198-4**
MC24-602-175, $^{1}$H NMR MeOD-$d_4$
Breaker 400MHz, 2024-06-25

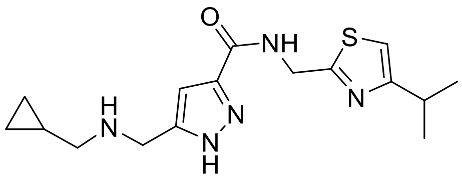


**5-(((Cyclopropylmethyl)amino)methyl)-N-((4-isopropylthiazol-2-yl)methyl)-1H-pyrazole-3-carboxamide (16).**

To a solution of **9.3** 5-formyl-N-((4-isopropylthiazol-2-yl)methyl)-1*H*-pyrazole-3-carboxamide (70 mg, 0.25 mmol) cyclopropylmethanamine (36 mg, 0.5 mmol) in MeOH (5 mL) was added AcOH (2 drops). The mixture was stirred at rt for 1h. The mixture was cooled to $0^{o}$C and $NaBH(OAc)_3$ (133 mg, 0.63 mmol) added. The mixture was stirred at rt for 5h, then diluted with EtOAc (10 mL) and water (5 mL). The organic phase was isolated, washed with brine, dried over $Na_2SO_4$, and concentrated to dryness. The crude material was purified by RP-HPLC to afford title compound **16** as a white solid (36 mg, 43%): LCMS [M+H] 334.2, ≥95% (214, 254 nm); HPLC ≥95% (254 nm); $^{1}$H NMR (400 MHz, MeOD) δ 8.53 (s, 1H), 7.05 (s, 1H), 6.93 (s, 1H), 4.81 (s, 2H), 4.27 (s, 2H), 3.13 – 2.98 (m, 1H), 2.90 (d, J = 7.4 Hz, 2H), 1.29 (d, J = 7.0 Hz, 6H), 1.17 – 1.00 (m, 1H), 0.80 – 0.60 (m, 2H), 0.47 – 0.28 (m, 2H).

### ***NMR***

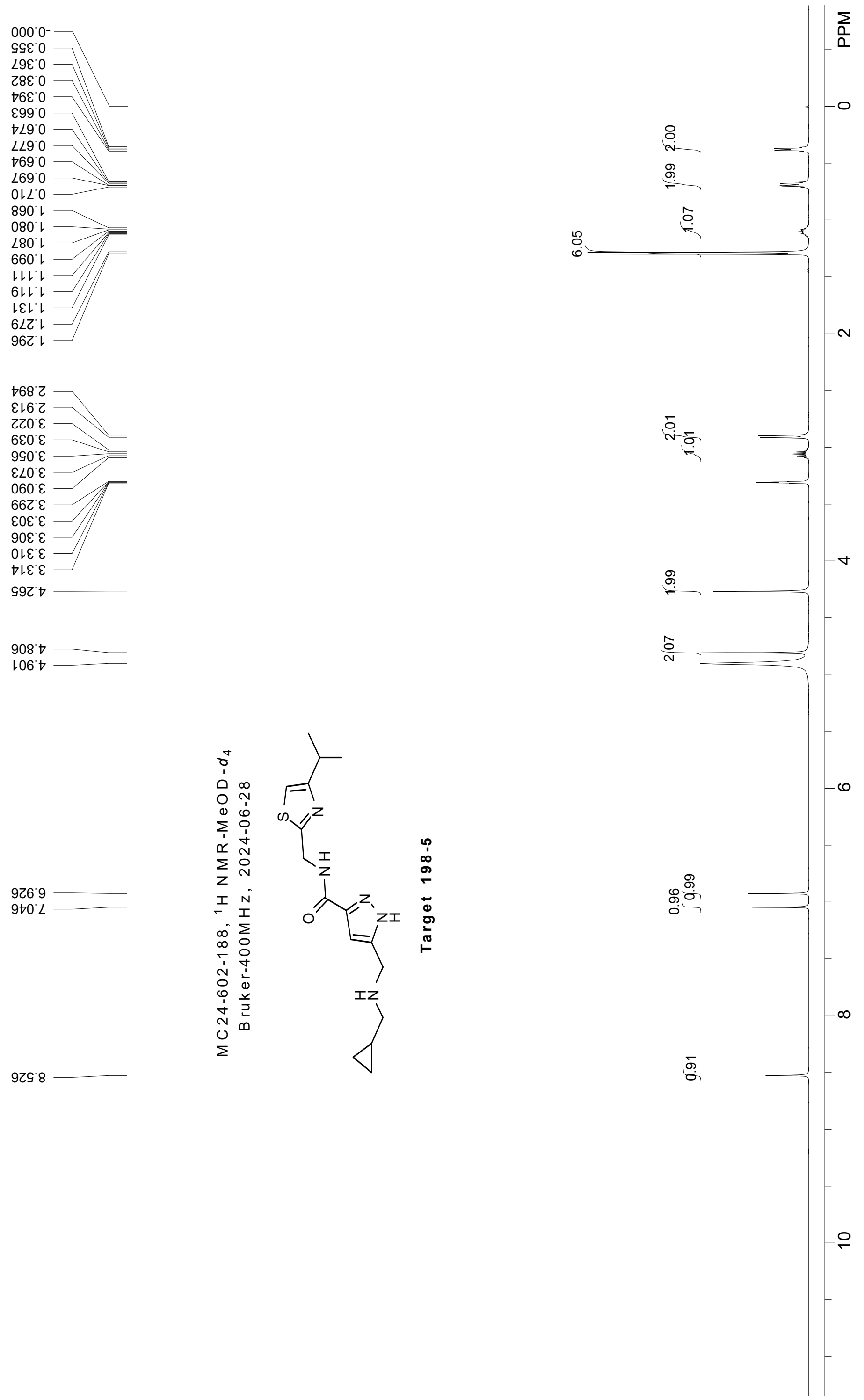
MC24-602-188, 1H NMR-MeOD-d4
Bruker-400MHz, 2024-06-28
Target 198-5
PPM
